# Neuroanatomical differences in early bilingual and monolingual children

**DOI:** 10.64898/2026.05.08.722956

**Authors:** Mia R. Coutinho, Guinevere F. Eden

## Abstract

Prior studies have reported inconsistent results for neuroanatomical differences between early bilinguals and monolinguals. These studies primarily measured gray matter volume (GMV), involved small samples, and prioritized adults. Few studies of early bilinguals have measured cortical thickness (CT), which offers more anatomical specificity. It remains unclear whether results derived from differing metrics and approaches (e.g., vertex-versus parcel-wise analyses) converge. Using data from the Adolescent Brain Cognitive Development^SM^ (ABCD) Study, we compared neuroanatomy between large groups of early cultural Spanish-English bilingual and English monolingual children (N = 1,209) matched on age, pubertal status, sex, handedness, socioeconomic status (SES), and nonverbal reasoning. Whole-brain voxel-based morphometry revealed areas of greater and of lesser GMV in bilinguals than monolinguals across all lobes. Vertex-wise CT analyses similarly identified widespread differences, with bilinguals showing areas of both thicker and thinner cortex. We contextualized these findings with parcel-wise CT analyses (average CT values), utilizing two atlases of differing spatial granularity. Parcel-wise results showed good correspondence with vertex-wise findings when implementing the more fine-grained atlas (Destrieux), but use of the coarser atlas (Desikan-Killiany) provided results that led to different conclusions. Finally, we tested for interaction effects between bilingualism and SES on CT and found several regions where differences between bilinguals and monolinguals in CT were modulated by SES. Together, these findings indicate that early bilingualism is associated with extensive neuroanatomical differences relative to monolinguals during childhood, and that these results can vary as a function of neuroanatomical metric, analysis approach, atlas granularity, and SES.

**Research Highlights:** Early Spanish–English bilingual and monolingual children differ in gray matter volume and cortical thickness across multiple brain regions.

Cortical thickness differences between bilinguals and monolinguals cannot be firmly attributed to adaptations associated with language or executive control.

Socioeconomic status modulates cortical differences between early bilinguals and monolinguals, revealing unique thickness patterns for those with lower versus higher SES backgrounds.

Parcel-wise between-group cortical thickness results are affected by atlas choice and can influence the interpretation of the findings.

## 1. Introduction

Bilingualism, or fluency in two languages, is estimated to be experienced by about half of the world’s population (Grosjean, 2021). Studies on bilinguals encompass a wide range of dual-language experiences, from those exposed to two languages from birth to others acquiring a second language proficiently later in life. The term “early bilingual” refers to those exposed to both languages during early childhood (often before or by age 7 years). In the United States, most of these individuals are “cultural early bilinguals” (also known as heritage speakers) because the language spoken at home is part of their family’s culture. Of the 21% of people who are bilingual in the United States, Spanish-English bilinguals are the most prevalent (United States Census Bureau, 2021). Here, we investigated differences in brain structure measured with magnetic resonance imaging (MRI) in early cultural bilingual children who speak Spanish and English to monolingual users of English. Confining the study to early bilinguals is important given that early bilinguals have been shown to differ from monolinguals in other ways than late bilinguals do (Claussenius-Kalman et al., 2020; Mechelli et al., 2004; Berken et al., 2017; Klein et al., 2019). The study of cultural early bilinguals provides a naturalistic way to determine the impact of an early dual-language experience on brain anatomy, thereby offering insights into the neuroscience of language and plasticity. The current study builds on a small corpus of work comparing brain structure between early bilinguals and monolinguals that has yielded mixed results. Further, it investigates two structural measures and two analysis approaches that have been employed separately across prior studies, thus providing a comprehensive picture of neuroanatomical differences in early bilinguals in a single study.

Considering comparison of brain structure between early bilinguals and monolinguals at the level of the whole brain, the first such study (Mechelli et al., 2004) found relatively greater gray matter density (GMD) in the left inferior parietal cortex in early bilingual adults and attributed this to the experience of acquiring a second language (Mechelli et al., 2004). Olulade et al., (2016) found relatively more gray matter volume (GMV) in early bilingual adults in bilateral frontal and parietal, left occipital, and right temporal areas. Based on these anatomical locations, some of these differences were attributed to executive control (EC), as EC has been proposed to be heightened in bilinguals due to a constant suppression and selection of one language over the other (Bialystok et al., 2012; Green & Abutalebi, 2013). They also reported less GMV in early bilinguals relative to monolinguals in right temporal, left occipital, and left and right cerebellar regions (Olulade et al., 2016). However, another GMV study did not find differences, although they did find greater GMV in left Heschl’s gyrus in the early bilingual adults when using a region-of-interest instead of the whole-brain analysis (Ressel et al., 2012). In children, García-Pentón and colleagues found more GMV in early bilinguals in a right parietal-occipital region, and Schug et al. (2022) found more GMV in bilateral frontal and right parietal areas in bilingual children and attributed these findings to EC differences. Schug et al. (2022) also reported less GMV in early bilingual relative to monolingual children in bilateral superior temporal and left inferior parietal cortices, the latter finding in direct opposition to the inferior parietal GMD result by Mechelli et al., (2004). Together, these studies offer little convergence in the location of the gray matter differences in bilinguals, as illustrated by no cortical findings in a meta-analysis that included most of these studies (Danylkiv & Krafnick, 2020). There are some investigations into cortical thickness (CT) in early bilinguals using a whole-brain, vertex-wise approach, and these have also conflicted in their results. In adults, Klein et al. (2014) found that early bilingual adults had thicker cortex in the left inferior frontal gyrus (IFG) and thinner cortex in the right IFG compared to monolinguals. Conversely, Nguyen et al., (2023) reported no CT differences for early bilingual children. Taken together, the findings on neuroanatomical differences in early bilinguals relative to monolinguals do not converge well across publications for either GMV or CT metrics.

The current study seeks to advance this work in early bilinguals by leveraging the Adolescent Brain Cognitive Development ^SM^ (ABCD) Study to surmount problems of insufficient statistical power due to small sample sizes that are likely to have yielded inaccurate results in many studies of bilinguals (Munson & Hernandez, 2019). The ABCD Study® consists of over 11,000 children representative of the United States population and has been used to compare large groups of bilinguals and monolinguals on cognitive tasks (Dick et al., 2019) and on neuroanatomical measures (Nguyen et al., 2024; Ronderos et al., 2024; Vaughn et al., 2021). The ABCD Study also affords an opportunity to focus on children, who are understudied in the general neuroimaging literature, due to the challenges that come with data acquisition and analysis of pediatric participants. However, if brain structure differs in bilingual children relative to their monolingual peers, bilinguals may need to be given special consideration in brain imaging studies. Importantly, it may be necessary to conduct brain imaging studies of common childhood conditions such as attention deficit hyperactivity disorder (ADHD) or developmental dyslexia in bilinguals separately from monolinguals should it turns out that the dual-language experience interacts with (mask or accentuate) brain structural differences associated with these conditions.

Neuroanatomical differences in bilinguals relative to monolinguals have often been interpreted within theories that account for how the bilingual brain supports two languages by adaptations in language and/or executive control systems (Bialystok et al., 2012; Green & Abutalebi, 2013; Hernandez & Li, 2007; Hernandez et al., 2018; Pliatsikas et al., 2020; Stocco et al., 2010). At the same time, others have taken the position that the structural differences in bilinguals are not related to any one specific skill, such as executive control; rather they are a consequence of the complex management of two languages leading to broad, but not universal, adaptations in the brain (Blanco-Elorrieta & Caramazza, 2021; Blanco-Elorrieta, 2025). In the current study on early bilinguals, we addressed this question by testing for associations between brain structure (within those regions that differed between early bilinguals and monolinguals) and performance on language and executive control measures. Findings of brain-behavioral associations, especially if bilinguals and monolinguals exhibit discrepancies in their performance of these skills (e.g., Patra et al., 2020), would support the notion a specific role of language and/or EC, while null result would speak for broader adaptations that are not tied to a specific skill. While there have been no investigations of brain-behavior correlational analyses in early bilinguals, several have studied groups consisting of a mix of early and late adult bilinguals. Mechelli et al., (2004) reported a positive relationship between GMD in left inferior parietal cortex and language skills (a combination of four language measures) using a region-of-interest (ROI) analysis; and Grogan et al., (2012) reported such a positive relationship between GMD in the inferior frontal cortex for language skills (lexical efficiency) using a whole-brain analysis, but no results for correlations in ROIs placed in bilateral inferior parietal and left superior temporal cortices. Grogan also repeated these analyses for GMV (modulated images), but had null results.

Further, Nguyen et al., (2023) reported negative relationships between CT in left and right superior frontal regions and language skills (a combination of two or three language measures) using a whole-brain approach. The correlations in these three studies were conducted with language performance measures in the second language. Here we tested for associations between brain structure and performance on an English language measure, as well as an EC measure in early bilinguals, expecting that if brain structural differences in early bilinguals reside in brain regions that subserve language or executive control, CT in some of these regions would demonstrate associations with these skills.

Consistent with the increasingly preferred use of CT over GMV, the primary metric in the current study is vertex-based CT. However, we also measured voxel-wise GMV in addition to CT to make a connection to the published GMV studies on early bilinguals noted above, and to contribute to the effort of providing multiple structural measures for the same participant groups (Claussenius-Kalman et al., 2020). Further, we extended the investigation by also taking a parcel-wise analysis approach, since parcel-based values are commonly reported when utilizing the ABCD Study data. Preprocessed parcellated values provide convenience and consistency among ABCD Study-based investigations (Hagler et al., 2019). However, it is not clear the degree to which results generated from average CT values from atlas-based parcels are comparable to those results derived from the vertex-based approach, especially since the use of an atlas with fewer, larger parcels for registration has been shown to negatively impact the fidelity of the results (Fürtjes et al., 2023). To address this issue, CT data generated from the vertex-wise between-group comparison were first converted to a parcellated format that was then submitted to between-group comparison. Second, we used the parcellated data provided by the ABCD Data Analysis, Informatics & Resource Center for a separate between-group comparison. The latter is the approach used by Vaughn et al., (2021) to compare early bilingual and monolingual children from the ABCD Study. They found widespread thinner cortex across all lobes in both hemispheres in parcels defined by the Desikan-Killiany atlas (Desikan et al., 2006). In the current study, parcel-wise analyses were conducted with two of the available atlases to establish the role of atlas choice, as the Destrieux (Destrieux et al., 2010) provides greater spatial resolution than the Desikan-Killiany atlas, with 148 versus 68 parcels, respectively. These additional analyses will provide insights into whether the between-group results from these approaches are interpreted similarly, providing an important contribution given the difficulties in achieving converging results in this field (Fürtjes et al., 2023; García-Pentón et al., 2019)

Lastly, socioeconomic status (SES), commonly defined by parental educational attainment or family income (McLoyd, 1998), has been shown to have a relationship with neuroanatomy (Noble, 2015), notably in regions that subserve language and EC. This raises the question whether SES and bilingualism interact when it comes to brain structure. This question was investigated by Brito and Noble (2018), who found several neuroanatomical differences between bilingual and monolingual adolescents to be more pronounced in a group with lower than a group with higher SES. We therefore matched the two groups on measures of SES in the analyses for the current study, but we also conducted a separate analysis to explicitly test for an interaction effect of Language Background (Bilingual vs. Monolingual) and SES Status (Lower vs. Higher) on CT in our group of early bilinguals.

In sum, the current study tested for neuroanatomical differences between large groups of early cultural Spanish-English and English-speaking monolingual children and went on to investigate relationships between CT and performance on language and EC skills in those regions found to differ between early bilinguals and monolinguals. Based on prior studies in bilinguals who acquired their second language later, we predicted that early bilinguals would exhibit a combination of relatively thicker and thinner cortex in regions known to subserve language and/or executive control, and that CT in these areas may be related to performance in these skills. In addition to the vertex-wise analysis, we conducted parcel-wise analyses to establish if results derived from vertex-versus parcel-based analyses lead to the same interpretation of the structural impact of bilingualism. Lastly, based on the known modulatory effect of SES on neuroanatomical differences in bilinguals versus monolinguals, we used a factorial design in a vertex-wise analysis, expecting more pronounced between-group differences for CT in those from lower-, relative to those from higher-SES backgrounds. Together, this comprehensive approach will characterize brain structural differences observed in early bilingual children across varied analysis approaches, and shed light on the mechanisms that bring about dual-language induced plasticity.

## 2. Methods

### 2.1 Participants

The participants in this study were drawn from the ABCD Study (Annual Release 5.0), which includes longitudinal data from 11,868 children from 21 sites who were between nine and ten years old at the first timepoint. Unless otherwise noted, all demographic data for this current study came from this baseline timepoint.

#### 2.1.1 Language background criteria

Bilinguals were identified using the Youth Acculturation Survey (YAS) modified from PhenX which gathers demographic information. We included those who responded both “Yes” to the question, “Besides English, do you speak or understand another language or dialect?” and those who went on to respond “Spanish” to the question “What ‘other’ language do you speak or understand besides English?” Next, we identified those who consistently used Spanish by including only those who responded to “What language do you speak with most of your family?” with 1) “Non-English Language all the time,” 2) “Non-English Language most of the time,” or 3) “Non-English Language and English equally.” To ensure our bilinguals were heritage speakers, we only included children whose parents responded “Yes” when asked “Do you consider the child Hispanic/Latino/Latina?” on the Parent Longitudinal Demographics Survey (PLDS) (which was collected at the one-year follow-up timepoint). From these criteria, 1,103 cultural Spanish-English bilingual children were identified.

Monolinguals were included if they responded “No” to the questions, “Besides English, do you speak or understand another language or dialect?” on the YAS and if their parents reported on the PLDS that the child’s native language was “English” with the question “What is your child’s native language? In other words, what was the first language predominantly spoken to your child by their parent or guardian after birth?” Additionally, English-speaking monolinguals were only included if the parent indicated their child had not attended a dual-language program (“Does your child attend a dual-language or language immersion program at their school? In other words, are several of their classes taught in a language other than English?”). This yielded 6,779 English monolingual children.

#### 2.1.2 Cognitive criteria

We included only those participants with scaled scores ≥ 4 on an automated version of the Matrix Reasoning subtest from the Wechsler Intelligence Test for Children-V (Wechsler, 2014) and ≥70 on the National Institutes of Health (NIH) Toolbox Picture Vocabulary Test (Gershon et al., 2013, 2014; Luciana et al., 2018) to exclude participants with an intellectual disability disorder (i.e., those with scores two standard deviations below the mean). As studies have shown neuroanatomical differences in reading disability (Eckert et al., 2016), we included only those with scores ≥85 on aloud single-word reading measured via the NIH Toolbox Oral Reading Recognition Task (TORRT; Gershon et al., 2013, 2014). These steps resulted in 853 bilinguals and 5,744 monolinguals.

#### 2.1.3 Medical, neurological, and mental health criteria

We included only those participants whose parents did not report a serious medical or neurological condition on the ABCD Parent Screener (“Has your child been diagnosed with any of the following serious medical or neurological conditions by a health professional?”) These exclusionary conditions included brain aneurysm, autism spectrum disorder, cerebral palsy, brain hemorrhage, brain hematoma, substance use disorder, schizophrenia, tumor, stroke, or ‘other’ serious medical conditions. Further, we included only participants who scored below the clinically significant range (<70) on the following Child Behavior Checklist items to screen for potentially confounding mental health conditions: depression, anxiety, somatic processing, oppositional disorder, conduct disorder, and ADHD. We also only included participants with no “Unspecified Schizophrenia Spectrum and Other Psychotic Disorder” (either in the past or at the time of study) from the Diagnostic and Statistical Manual of Mental Disorders, Fifth Edition (DSM-5) Kiddie Schedule for Affective Disorders and Schizophrenia Present and Lifetime Version (KSADS-PL). Following these steps, 780 bilinguals and 5,163 monolinguals remained.

#### 2.1.4 ABCD Quality Control MRI Clinical Report

Participants were removed if the Quality Control MRI Clinical Report provided by the ABCD Study indicated MR images of poor quality (“image artifacts prevent radiology read”) or brain abnormalities (“consider clinical referral” or “consider immediate clinical referral”). We included only those with “no abnormal findings” or “normal anatomical variant of no clinical significance” (rating 1 and 2). Following this, 756 bilinguals and 4,598 monolinguals remained.

#### 2.1.5 Group Matching

The bilingual and monolingual participants were matched on age, pubertal status, sex, handedness, SES (both household income and parental education), and nonverbal reasoning. Propensity matching was conducted using the PsmPy package (Kline & Luo, 2022). Age was based on months; pubertal status, on the Parent Pubertal Development Scale (Barch et al., 2018; Petersen et al., 1988), and handedness, on a brief version of Edinburgh Handedness Inventory (Oldfield, 1971; Veale, 2014). SES for household income and parental education were based on the Parent Longitudinal Demographics Survey. Household income bins were converted to income amount and ranged from $0 to $300,000; parental education ranged from 0 to 21 (see Table 1 caption, below). Following this, 619 early cultural Spanish-English bilinguals and 640 monolinguals remained.

**Table 1.** Demographic and performance measures for the Bilingual and Monolingual Groups.

|  | **Bilinguals** | **Monolinguals** | **Statistics: Student's t-test or chi-square** |
| --- | --- | --- | --- |
| N | 601 | 608 | - |
| Age (years) | 9.82 ± 0.62 | 9.83 ± 0.60 | n.s. |
| Pubertal Status - Female | 1.00 ± 1.25 | 1.10 ± 1.26 | n.s. |
| Pubertal Status - Male | 0.80 ± 0.84 | 0.77 ± 0.94 | n.s. |
| Sex (M/F) | 285/316 | 295/313 | n.s. |
| Handedness (R/Non-R) | 508/93 | 489 /119 | n.s. |
| SES: Household Income (USD) | 54,360 ± 49,710 | 57,500 ± 58,065 | n.s. |
| SES: Parental Education | 14.82 ± 3.43 | 15.15 ± 2.52 | n.s. |
| Nonverbal Reasoning (MR) | 285/316 | 295/313 | n.s. |
| Vocabulary (PVT) | 100.88 ± 13.80 | 104.50 ± 15.38 | p<0.05 |
| Language Learning (RAVLT) | 6.34 ± 2.69 | 5.95 ± 2.65 | p<0.05 |
| Inhibitory Control (Flanker) | 95.33 ± 12.77 | 95.83 ± 14.65 | n.s. |
| Visuospatial (Little Man) | 0.59 ± 0.16 | 0.57 ± 0.17 | p<0.05 |

*Note:* Counts reported for group size, sex, and handedness. Averages and standard deviation reported for age, pubertal status, SES scores, nonverbal reasoning, vocabulary, language learning, inhibitory control, and visuospatial processing. Statistical tests: Student's t-test or chi-square, are reported in the right column. Pubertal status=Pubertal developmental scale; Sex: M=Male, F=Female; Handedness=Brief Edinburgh Handedness. SES Household Income=Parent Longitudinal Demographics Survey, in which household income bin was converted to income amount; SES Parental Education=Binned variable in which 14=General Education Development (GED) Diploma or equivalent, and 15=Some college. Nonverbal Reasoning= Matrix Reasoning (MR). Vocabulary=NIH Toolbox Picture Vocabulary Test (PVT) age-corrected standard score; Language Learning=Rey Auditory Verbal Learning Test (RAVLT) Learning score (Trial V-Trial I); Inhibitory Control=NIH Toolbox Flanker Task accuracy; Visuospatial Processing= Little Man Task percent correct. (n.s.=not significant, $p \geq 0.05$ )

**Table 2.** Location of cortical thickness differences between the Bilingual and Monolingual Groups (vertex-wise analysis).

| MNI Coordinates |  |  |  |  |  |  |
| --- | --- | --- | --- | --- | --- | --- |
| x | y | z | Region | BA | Z | K<sub>E</sub> |
| **Bilinguals > Monolinguals** |  |  |  |  |  |  |
| **-33** | **26** | **-9** | **Left Inferior Orbitofrontal Gyrus** | **47** | **4.62** | **202** |
| -32 | 19 | -22 | Left Inferior Orbitofrontal Gyrus | 47 | 3.11 |  |
| -33 | 9 | 12 | Left Insula | 13 | 2.85 |  |
| **-6** | **24** | **-24** | **Left Gyrus Rectus** | **11** | **4.15** | **109** |
| -10 | 16 | -17 | Left Olfactory Gyrus | 11 | 4.05 |  |
| **-21** | **-25** | **-20** | **Left Parahippocampal Gyrus** | **36** | **4.45** | **105** |
| **-22** | **-100** | **-13** | **Left Lingual Gyrus** | **18** | **7.52** | **607** |
| -29 | -91 | 8 | Left Middle Occipital Gyrus | 18 | 3.81 |  |
| **47** | **-17** | **3** | **Right Superior Temporal Gyrus** | **41** | **3.88** | **152** |
| 59 | -9 | 2 | Right Superior Temporal Gyrus | 41 | 3.54 |  |
| 56 | -2 | -3 | Right Superior Temporal Gyrus | 22 | 3.02 |  |
| **24** | **-100** | **-6** | **Right Inferior Occipital Gyrus** | **18** | **7.84** | **359** |
| **Monolinguals > Bilinguals** |  |  |  |  |  |  |
| **-43** | **26** | **27** | **Left Inferior Frontal Gyrus pars triangularis** | **9** | **3.82** | **162** |
| **-31** | **12** | **49** | **Left Middle Frontal Gyrus** | **6** | **4.37** | **386** |
| -21 | 9 | 57 | Left Superior Frontal Gyrus | 6 | 4.05 |  |
| -23 | 22 | 49 | Left Middle Frontal Gyrus | 8 | 3.33 |  |
| **-8** | **65** | **5** | **Left Medial Superior Frontal Gyrus** | **10** | **4.71** | **168** |
| -4 | 60 | -14 | Left Gyrus Rectus | 10 | 3.57 |  |
| **-6** | **8** | **60** | **Left Supplementary Motor Area** | **6** | **4.92** | **391** |
| -7 | 36 | 42 | Left Medial Superior Frontal Gyrus | 8 | 4.42 |  |
| -6 | -15 | 61 | Left Supplementary Motor Area | 6 | 4.18 |  |
| **-20** | **-17** | **67** | **Left Precentral Gyrus** | **6** | **3.93** | **419** |
| -24 | -31 | 58 | Left Postcentral Gyrus | 1 | 3.42 |  |
| -11 | -25 | 75 | Left Paracentral Lobule | 4 | 3.25 |  |
| **-53** | **-15** | **36** | **Left Postcentral Gyrus** | **1** | **4.55** | **658** |
| -34 | -20 | 39 | Left Postcentral Gyrus | 1 | 3.41 |  |
| -60 | -8 | 16 | Left Postcentral Gyrus | 1 | 3.32 |  |
| **-52** | **-21** | **20** | **Left Supramarginal Gyrus** | **40** | **4.72** | **313** |
| -45 | -13 | 17 | Left Rolandic Opercularis | 4 | 3.56 |  |
| **-35** | **-33** | **-20** | **Left Fusiform Gyrus** | **36** | **3.8** | **126** |
| -38 | -21 | -26 | Left Fusiform Gyrus | 36 | 3.66 |  |
| -31 | -44 | -15 | Left Fusiform Gyrus | 37 | 3.12 |  |
| **-7** | **-64** | **54** | **Left Precuneus** | **7** | **3.68** | **117** |
| -12 | -75 | 49 | Left Precuneus | 7 | 3.3 |  |
| **-10** | **-67** | **27** | **Left Cuneus** | **7** | **5.06** | **177** |
| -3 | -63 | 19 | Left Calcarine Fissure | 31 | 3.34 |  |
| -7 | -57 | 23 | Left Precuneus | 31 | 2.74 |  |
| **-14** | **-73** | **18** | **Left Calcarine Fissure** | **17** | **6.11** | **852** |
| -11 | -83 | 3 | Left Calcarine Fissure | 17 | 5.32 |  |
| -4 | -79 | 25 | Left Cuneus | 18 | 4.82 |  |
| -3 | -79 | 14 | Left Calcarine Fissure | 18 | 4.41 |  |
| **-48** | **15** | **-23** | **Left Superior Temporal Pole** | **38** | **5.69** | **155** |
| **8** | **58** | **-3** | **Right Medial Orbitofrontal Gyrus** | **10** | **3.65** | **188** |
| 16 | 63 | -6 | Right Superior Orbitofrontal Gyrus | 10 | 3.28 |  |
| 24 | 64 | 2 | Right Superior Frontal Gyrus | 10 | 3.09 |  |
| **6** | **7** | **60** | **Right Supplementary Motor Area** | **6** | **4.35** | **348** |
| 7 | 22 | 49 | Right Supplementary Motor Area | 8 | 4.04 |  |
| 5 | -16 | 57 | Right Supplementary Motor Area | 6 | 3.6 |  |
| **35** | **-20** | **43** | **Right Precentral Gyrus** | **4** | **4.14** | **421** |
| 53 | -7 | 22 | Right Rolandic Opercularis | 4 | 3.91 |  |
| 57 | -10 | 34 | Right Postcentral Gyrus | 4 | 3.39 |  |
| **21** | **-27** | **55** | **Right Postcentral Gyrus** | **1** | **3.94** | **499** |
| 11 | -34 | 64 | Right Postcentral Gyrus | 4 | 3.67 |  |
| 7 | -39 | 58 | Right Precuneus | 5 | 3.3 |  |
| **3** | **-62** | **26** | **Right Precuneus** | **31** | **4.48** | **271** |
| 4 | -67 | 39 | Right Precuneus | 7 | 3.2 |  |
| **3** | **-72** | **20** | **Right Cuneus** | **18** | **4.64** | **180** |
| **14** | **-74** | **-6** | **Right Calcarine Fissure** | **18** | **3.6** | **151** |
| 5 | -75 | 6 | Right Calcarine Fissure | 17 | 3.36 |  |
| **41** | **-15** | **21** | **Right Rolandic Opercularis** | **1** | **3.55** | **147** |
| 37 | -18 | 15 | Right Insula | 13 | 3.24 |  |
| **49** | **11** | **-17** | **Right Superior Temporal Pole** | **38** | **4.79** | **201** |
| 55 | 3 | -19 | Right Middle Temporal Gyrus | 38 | 3.22 |  |
| 38 | 13 | -40 | Right Middle Temporal Pole | 38 | 3.04 |  |

Note: Reported using MNI coordinates. Bold font indicates the peak coordinate of a cluster while gray font indicates the subpeaks within clusters.

### 2.2 MRI data

#### 2.2.1 Acquisition

Images for all participants (downloaded from the ABCD Study Data Repository) consisted of T1-weighted MR images acquired on 3T MR scanners with a 1×1×1 mm^3^ voxel resolution. Details on acquisition protocols can be found at https://abcdstudy.org/images/Protocol_Imaging_Sequences.pdf.

#### 2.2.2 MRI CAT12 quality control

After completing image segmentation using the Computational Anatomy Toolbox 12 software (CAT12), quality control was performed using the CAT12 Weighted Overall Image Quality assessment. The resulting image quality ratings (IQR values) that were calculated ranged from 0.5 (excellent) to 10.5 (unacceptable) and then were converted to percentage ratings based on the CAT12 manual (for more information, see the CAT12 manual: https://neuro-jena.github.io/cat12-help/). Images that scored below 75% on this quality assessment were excluded. Next, participant images were visually inspected by two researchers blinded to the groups to identify and remove those with skull-stripping/segmentation or file corruption errors.

Following these steps, the final sample was 601 bilinguals and 608 monolinguals. The groups (see Table 1, below), continued to be matched on age, pubertal status, sex, handedness, SES (household income and parental education), and nonverbal reasoning. Despite excluding those with ADHD, Child Behavioral Checklist scores for ADHD were higher in the Monolingual than Bilingual Group and were therefore added as a covariate of no interest in the MRI data analyses.

### 2.3 MRI Data Analyses

#### 2.3.1 Bilinguals versus monolinguals voxel-wise gray matter volume analysis

First, to contextualize our results within the GMV literature on early bilinguals, we compared the early bilinguals with monolinguals on GMV using the automated Voxel-Based Morphometry technique (Ashburner & Friston, 2000) in CAT12 (Gaser et al., 2022) run through in the statistical parametric mapping (SPM) 12 software (Dahnke et al., 2013) utilizing the Template-O-Matic (TOM8) Toolbox to generate an age- and sex-specific tissue probability map. Preprocessing steps included segmentation, normalization, and modulation to adjust for total intracranial volume. A between-group t-test was performed to compare the Bilingual (n=601) and Monolingual (n=608) Groups on whole-brain GMV with the following covariates of no interest: age, pubertal status, sex, SES (household income and parent education), nonverbal reasoning, ADHD, scanner site, scanner model and total GMV. Statistical thresholds were a voxel-wise height threshold of p <0.005 (uncorrected) and a cluster-level extent threshold of p<0.05 (FDR corrected).

#### 2.3.2 Bilinguals versus monolinguals vertex-wise cortical thickness analysis

For our main goal of comparing CT between bilinguals and monolinguals, we used the CAT12 Toolbox (Gaser et al., 2022) following the CAT12 Manual (Dahnke et al., 2013). The images were preprocessed as described in Section 2.3.1. A between-group t-test for CT was performed on the same 601 bilingual and 608 monolingual participants as conducted for GMV. Covariates of no interest included age, pubertal status, sex, SES (household income and parent education), nonverbal reasoning, ADHD, scanner site and scanner model. Statistical thresholds were identical to those described in Section 2.3.1.

For both GMV and CT, the coordinates of all resultant clusters were entered into the label4MRI package (https://github.com/yunshiuan/label4MRI), which uses the Atlas of Brodmann’s Areas and the Automated Anatomical Labeling Atlas (Tzourio-Mazoyer et al., 2002) to identify the corresponding Brodmann’s Areas and anatomical brain region for each cluster.

#### 2.3.3 Correlations between cortical thickness and skill performance in regions that differ between bilinguals and monolinguals

The regions that emerged from the bilingual versus monolingual CT comparison were tested for associations with performance in the domains of language and executive control skills (in the two groups combined). The specific skills were chosen based on known performance differences between bilinguals and monolinguals. Language measures included vocabulary measured via the NIH Toolbox Picture Vocabulary Task (PVT, noted above), and an automated version of the Rey Auditory Verbal Learning Test (RAVLT) Learning Score (Spreen & Strauss, 1998). Executive control was measured with the NIH Toolbox Flanker Task, a variant of the Eriksen Flanker Task (Eriksen & Eriksen, 1974), to gauge inhibitory control. Additionally, we tested for associations with visuospatial processing using the Little Man Task (Acker & Acker, 1982) selected post-hoc following CT difference findings in occipital cortex; it simultaneously served as a cognitive measure not emphasizing language or EC.

The NIH Toolbox PVT requires participants to select the picture that best corresponded to a spoken word. The automated RAVLT involves reading a list of 15 words to a participant and asking them to recall the words from memory for five consecutive trials. The “RAVLT Learning” score is the difference between Trials V and I. The NIH Toolbox Flanker Task requires participants to identify the direction of a central arrow while ignoring incongruent surrounding (flanking) arrows. The Little Man Task requires participants to view various images of a figure (‘little man’) holding a suitcase presented in different orientations and use mental rotation skills to assess which hand (left or right) is holding the suitcase. For further detail about these measures see Luciana et al., (2018). Performance scores were compared between Bilinguals and Monolinguals using a student’s t-test (p < 0.05; see Table 1).

To test for associations between CT (in those regions where bilinguals differed from monolinguals) and these cognitive skills, we created two masks in SPM12, one for each contrast (Bilingual>Monolingual Groups; Monolingual>Bilingual Groups) and performed a regression analysis of CT and the four performance measures restricted to these masked areas for all participants. Covariates of no interest were the same as those used for the Bilingual versus Monolingual Group vertex-wise CT analysis described above (Section 2.3.2).

#### 2.3.4 Bilinguals versus monolinguals parcel-wise cortical thickness analysis

Next, we extended the comparison of Bilinguals and Monolinguals on CT from the vertex-wise approach (Section 2.3.2) to the parcel-wise approach. We conducted two parcel-wise analyses, the first based on calculating parcel-based average CT values from our vertex-wise analysis, and the second based on the ABCD-processed parcel-based average CT values. For each, we used parcels registered to the Destrieux atlas (Destrieux et al., 2010), as well as to the Desikan-Killiany atlas (Desikan et al., 2006) to attain an understanding of the role of these two atlases’ distinct spatial resolution on the results.

##### 2.3.4.1 Parcel-wise analyses using CT values derived from vertex-wise analysis

We used the “ROI Tools” option in CAT12 (Gaser et al., 2022) to calculate and extract each participant’s average CT from each parcel for both atlases (Destrieux et al., 2010; Desikan et al., 2006).

##### 2.3.4.2 Parcel-wise analyses using CT values from ABCD Data Analysis, Informatics & Resource Center

We used the parcellated data provided by the ABCD Data Analysis, Informatics & Resource Center (Hagler et al., 2019) for all participants’ average CT within each parcel for both atlases (Destrieux et al., 2010; Desikan et al., 2006).

For both parcel-wise analyses, one-way, two-sided, between-group ANCOVAs were used for each parcel to compare CT between the Bilingual and the Monolingual Groups (68 ANCOVAs for the Desikan-Killiany, and 148 for the Destrieux atlas). Statistical analyses were conducted in Python using a statistical threshold of p<0.05, FDR-corrected, and the same covariates of no interest as those in the vertex-wise analysis (Section 2.3.2). Statistical values for all parcels were displayed as a surface map using the *ggseg* R package (Mowinckel & Vidal-Piñeiro, 2020).

#### 2.3.5 ANOVA to test for an interaction of Language Background (Bilingual vs. Monolingual) and SES (Lower vs. Higher)

##### 2.3.5.1 Selection of the Bilingual and Monolingual Groups of Lower and Higher SES

From the sample used above, Lower-SES and Higher-SES Groups were created by using the converted score (from a binned value ranging from 1 to 10, to the midpoint value of each bin, e.g., Bin 1=less than $5,000: converted value =$2,500; Bin 9*=$100,000 through $199,999: converted value=$150,000; Bin 10=$200,000 and greater;* converted value= $300,000) to divide the groups by median household income for the full sample ($54,208). We then matched bilinguals with monolinguals for the two Lower-SES Groups, as well as the two Higher-SES Groups on both measures of SES (household income and parental education). This resulted in 322 participants in the Bilingual Lower-SES Group, 318 in the Monolingual Lower-SES Group, 223 in the Bilingual Higher-SES Group, and 247 in the Monolingual Higher-SES Group (see Table 3).

##### 2.3.5.2 2 × 2 ANOVA to test for interaction of Language Background (Bilingual vs. Monolingual) and SES (Lower vs. Higher) on CT

Using the vertex-wise approach, we conducted a 2 × 2 ANOVA to test for an interaction of Language Background (Bilingual vs. Monolingual) and SES (Lower vs. Higher) on CT. Covariates of no interest were the same as those used for the bilinguals versus monolinguals CT analysis above (Section 2.3.2), except for the SES measures. We applied the same vertex-wise height threshold of p<0.005, FDR; cluster-level extent threshold of p<0.05. To determine the direction of the effect for significant interactions, we extracted mean CT for each of the clusters that showed a significant Language Background by SES interaction on the F-map and conducted Student’s t-tests (statistical threshold = p<0.05).

## 3. Results

### 3.1 Bilinguals and monolinguals demographic and performance measures

Demographic and performance measures are provided in Table 1. The Bilinguals and Monolingual Groups were matched on age, pubertal status, sex, handedness, SES (household income and parental education) and nonverbal reasoning. For language performance, the Bilingual Group had a smaller vocabulary in English than the Monolingual Group did, as is typical, while they scored higher on language learning than the Monolingual

Group. For EC, there were no differences between the two groups on inhibitory control. Lastly, the bilinguals performed relatively better on visuospatial processing.

### 3.1 Bilinguals versus monolinguals voxel-wise gray matter volume results

As shown in Figure 2 and Supplemental Table 1, bilinguals exhibited more GMV than monolingual, (shown in red) in four left hemisphere and three right hemisphere clusters. Left hemisphere cluster peaks were located in (1) supplementary motor area (BA 6), with subpeaks extending to superior frontal gyrus (BA 6) and supplementary motor area (BA 6); (2) middle temporal gyrus (BA 21); (3) postcentral gyrus (BA 1) extending to the precuneus (BA 7) and superior parietal lobule (BA 7); and (4) calcarine fissure (BA 18) extending to inferior occipital gyrus (BA 18) and middle occipital gyrus (BA 18). In the right hemisphere, cluster peaks were in (1) olfactory gyrus (BA 11) extending to hippocampus (BA 28) and lingual gyrus (BA 18); (2) postcentral gyrus (BA 1) extending to the precuneus (BA 31, BA 7); and (3) calcarine fissure (BA 18) extending to the inferior occipital (BA 18) and middle occipital (BA 18) gyri.

**Figure 1:**
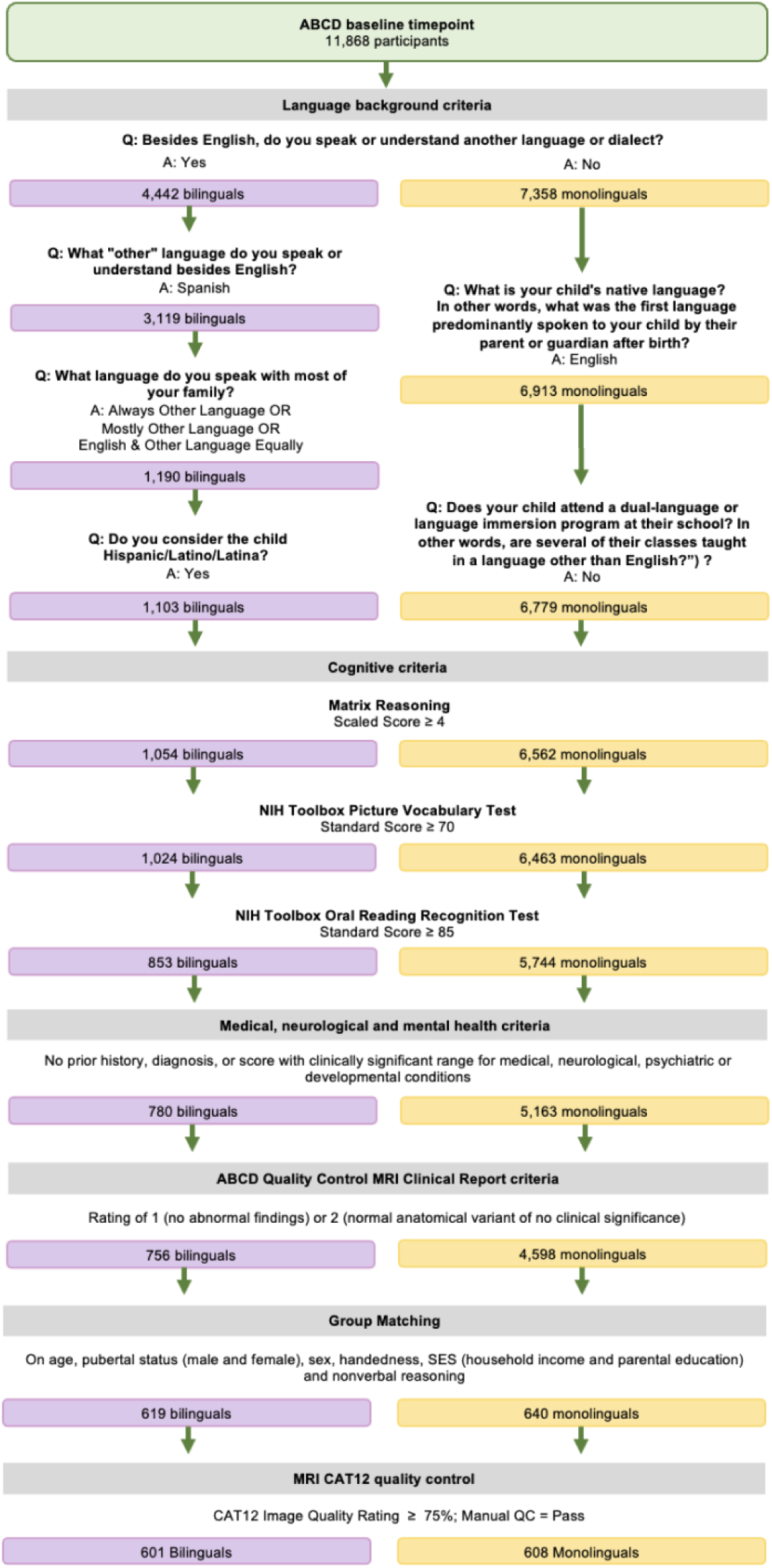
Selection of the Bilingual group (purple) and the Monolingual group (yellow) from the ABCD Study. *Note:* More information on the ABCD Study is outlined in Garavan et al., (2018). Participants with missing values for any variable used in the participant selection pipeline were excluded.

**Figure 2.**
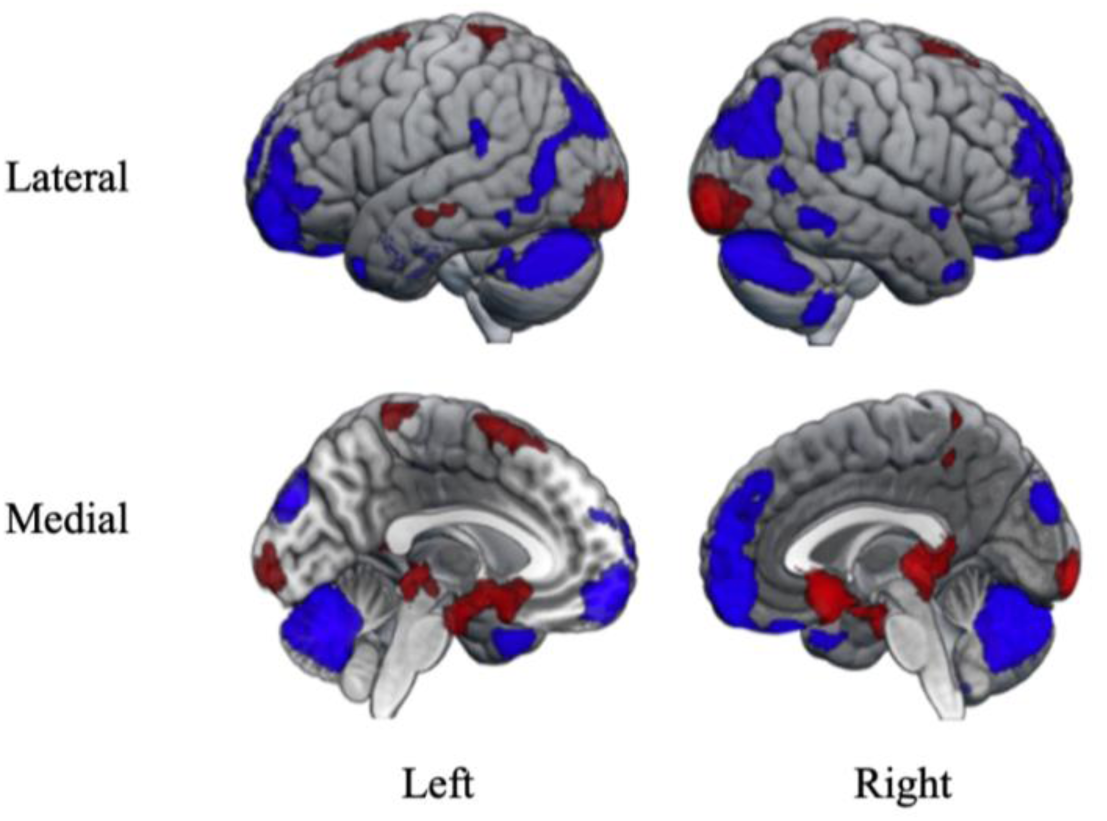
Voxel-wise gray matter volume differences between the Bilingual and Monolingual Groups. *Note*: Clusters with more GMV in bilinguals than monolinguals (red) and less GMV in bilinguals than monolinguals (blue). Voxel-wise height threshold p<0.001; FDR cluster-level extent threshold p<0.05. Coordinates of peaks and subpeaks for each cluster are found in Supplementary Table 1.

For the opposite contrast, bilinguals showed less GMV than monolinguals in five clusters in the left hemisphere and four clusters in the right hemisphere (shown in blue in Figure 2 and listed in Supplemental Table 1). In the left hemisphere, cluster peaks were located in (1) the medial superior frontal gyrus (BA 10), extending to the gyrus rectus (BA 11); (2) the superior temporal pole (BA 38), extending to additional subpeaks in the same region (BA 38) and to the middle temporal pole (BA 38); (3) the supramarginal gyrus (BA 40), extending to the superior temporal gyrus (BA 40, 22); (4) the middle occipital gyrus (BA 19), extending to the cuneus (BA 19); and (5) the cerebellum Crus I, extending to vermis VI and bilaterally to Crus 1 on the right. In the right hemisphere, cluster peaks were located in (1) the superior temporal gyrus (BA 22), extending to the angular gyrus (BA 39); (2) the fusiform gyrus (BA 37), extending to the inferior temporal gyrus (BA 37); (3) the middle occipital gyrus (BA 19); and (4) cerebellar lobule VIII, extending to cerebellar lobule IX.

### 3.2 Bilinguals versus monolinguals vertex-wise cortical thickness results

As shown in Figure 3 and Table 3, bilinguals exhibited thicker cortices compared to monolinguals in four left and two right hemisphere regions. On the left, cluster peaks were located in (1) inferior orbitofrontal gyrus (BA 47), extending into a second cluster in this gyrus (BA 47) and to the insula (BA 13); (2) gyrus rectus (BA 11), extending to olfactory gyrus (BA 11); (3) the parahippocampal gyrus (BA 36) and (4) the lingual gyrus (BA 18), extending to middle occipital gyrus (BA 18). On the right, peaks were in (1) superior temporal gyrus (BA 41) with two further subpeaks in this gyrus (BA 41, 22), and (2) inferior occipital gyrus (BA 18).

**Figure 3.**
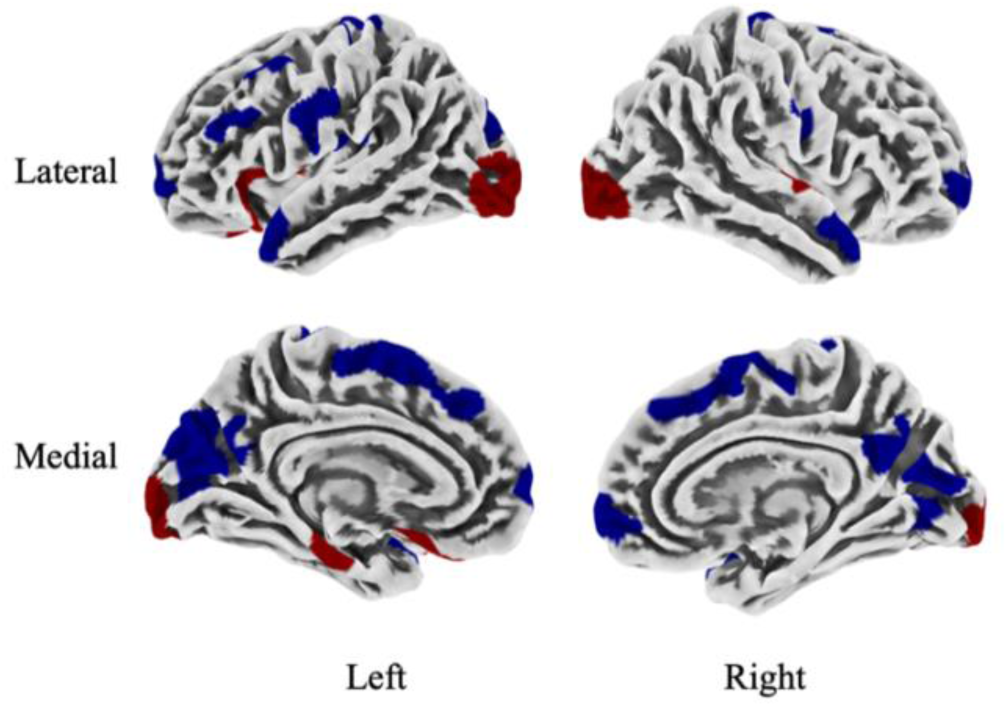
Vertex-wise cortical thickness differences between the Bilingual and Monolingual Groups. Note: Clusters with thicker cortex in bilinguals than monolinguals (red) and thinner cortex in bilinguals than monolinguals (blue). Vertex-wise height threshold p<0.005; FDR cluster-level extent threshold p<0.05. Coordinates of peaks and subpeaks for each cluster are found in Table 4.

The opposite contrast revealed 12 left and nine right hemisphere clusters where bilinguals exhibited thinner cortices compared to monolinguals. The 12 left hemisphere peaks were located in (1) inferior frontal gyrus *pars triangularis* (BA 9); (2) middle frontal gyrus (BA 6) extending to superior frontal gyrus (BA 6) and middle frontal gyrus (BA 8); (3) medial superior frontal gyrus (BA 10) extending to gyrus rectus (BA 10); (4) supplementary motor area (BA 6) extending to medial superior frontal gyrus (BA 8) and supplementary motor area (BA 6); (5) precentral gyrus (BA 6) extending to postcentral gyrus (BA 1) and paracentral lobule (BA 4); (6) postcentral gyrus (BA 1) extending to two further subpeaks in the same gyrus; (7) supramarginal gyrus (BA 40) extending to rolandic opercularis (BA 4); (8) the fusiform gyrus (BA 36) extending to two further subpeaks in the same gyrus (BA 36 and 37); (9) the precuneus (BA 7) extending to precuneus (BA 7); (10) the cuneus (BA 7) extending to calcarine fissure (BA 31) and precuneus (BA 31); (11) the calcarine fissure (BA 17) extending to a subpeak in calcarine fissure (BA 17,18), and to cuneus (BA 18); and (12) to superior temporal pole (BA 38). The nine clusters in the right hemisphere were located in (1) medial orbitofrontal gyrus (BA 10) extending to superior orbitofrontal gyrus (BA 10) and superior frontal gyrus (BA 10); (2) supplementary motor area (BA 6) extending to two further subpeaks in the same area (BA 6,8); (3) precentral gyrus (BA 4) extending to rolandic opercularis (BA 4) and postcentral gyrus (BA 4); (4) postcentral gyrus (BA 1) extending to a further subpeak within the same gyrus (BA 4) and to precuneus (BA 5); (5) precuneus (BA 31) extending to two further subpeaks in the same region (BA 7, 5); (6) cuneus (BA 18); (7) calcarine fissure (BA 18) extending to another subpeak in the region (BA 17); (8) rolandic opercularis (BA 1) extending to insula (BA 13); (9) superior temporal pole (BA 38) extending to middle temporal gyrus (BA 38) and middle temporal pole (BA 38).

### 3.3 Correlations between cortical thickness and skill performance in regions that differ between bilinguals and monolinguals

No significant correlations were observed between CT and task performance on either vocabulary, language learning, inhibitory control, or visuospatial processing (in regions where CT was found to differ between the Bilingual and Monolingual Groups).

### 3.4. Bilinguals versus monolinguals parcel-wise cortical thickness results

#### 3.4.1 Parcel-wise CT values derived from vertex-wise analysis

For the Destrieux atlas, bilinguals exhibited thicker cortex than monolinguals in five (out of 74) left hemisphere parcels (spanning frontal, temporal, and occipital regions) and three (out of 74) right hemisphere parcels (temporal and occipital regions). For the reverse comparison, bilinguals exhibited thinner cortex than monolinguals in 11 left (frontal, parietal, occipital), and seven (frontal, parietal, occipital) right-hemisphere parcels (Figure 4 and Supplemental Table 2).

**Figure 4.**
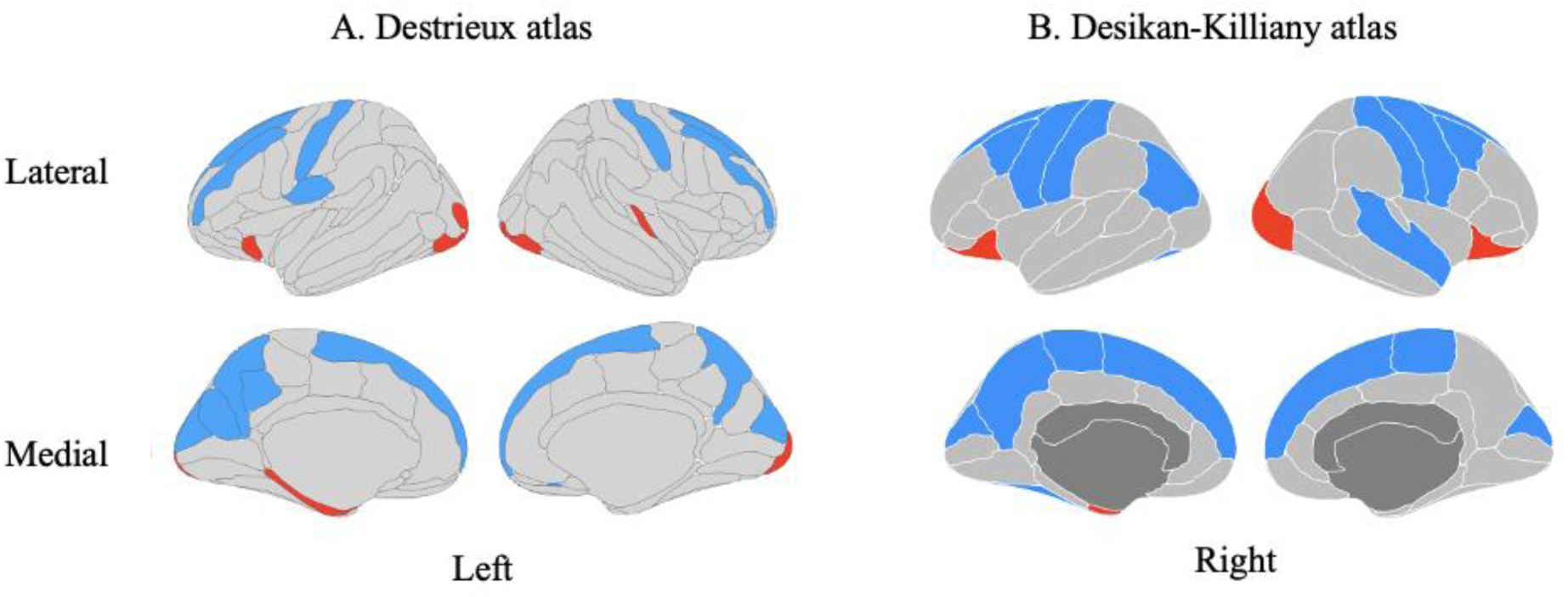
Parcel-wise cortical thickness differences between the Bilinguals and Monolinguals Groups (using vertex-wise analyses-derived values). *Note:* Parcels with greater CT in bilinguals than monolinguals (red) and less CT in bilinguals than monolingual (blue). Results were generated by registering data to the (A) Destrieux and (B) Desikan-Killiany atlases. Between-group ANOVA (p<0.05 uncorrected, FDR corrected). Results are listed in Supplemental Tables 2 and 3.

Using the Desikan-Killiany atlas, bilinguals exhibited thicker cortices compared to monolinguals in two (out of 34) left (frontal and temporal) and two (out of 34) right (frontal and occipital) hemisphere parcels. For the reverse comparison, bilinguals exhibited thinner cortex than monolinguals in nine left (frontal, parietal, temporal, occipital), and seven (frontal, temporal, parietal, occipital) right-hemisphere parcels (Figure 4 and Supplemental Table 3).

#### 3.4.2 Parcel-wise CT values from the ABCD Data Analysis, Informatics & Resource Center

For the same participants, when using ABCD-provided average CT values and the Destrieux atlas, bilinguals exhibited 21 (out of 74) parcels of thicker cortex in the left hemisphere (frontal, parietal, temporal, occipital) compared to monolinguals, and 24 (out of 74) parcels of thicker cortex in the right hemisphere (frontal, parietal, temporal, occipital). For the reverse comparison, bilinguals exhibited 36 parcels of thinner cortex in the left hemisphere (frontal, parietal, temporal, occipital), and 37 parcels of thinner cortex in the right hemisphere compared to their monolingual peers (frontal, parietal, temporal, occipital) (Figure 5 and Supplemental Table 4).

**Figure 5.**
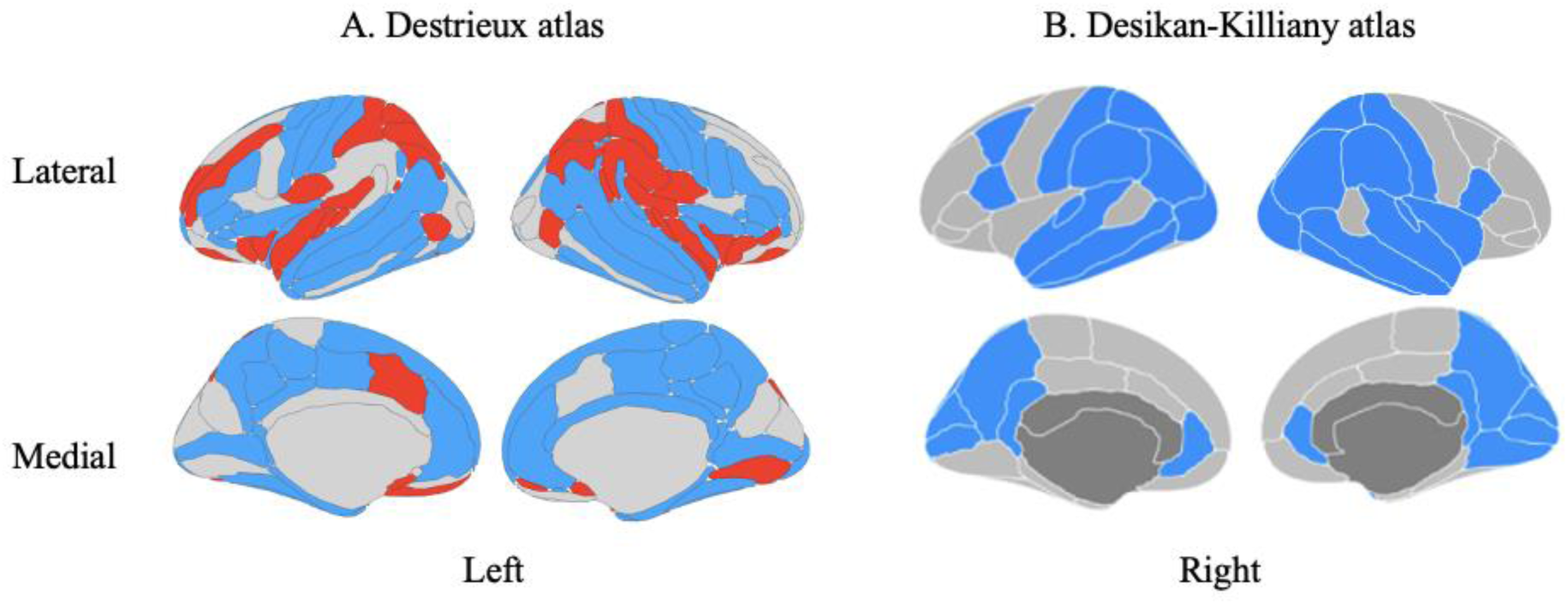
Parcel-wise cortical thickness differences between the Bilingual and Monolingual Groups (using ABCD-derived values). *Note:* Parcels with greater CT in bilinguals than monolinguals (red) and less CT in bilinguals than monolinguals (blue). Results were generated by registering data to the (A) Destrieux and (B) Desikan-Killiany atlases. Between-group ANOVA (p< .05 uncorrected, FDR corrected). Results are listed in Supplemental Tables 4 and 5.

Using the Desikan-Killiany atlas, bilinguals exhibited no parcels of significantly thicker cortices compared to monolinguals in either hemisphere. However, bilinguals exhibited 17 (out of 34) parcels of thinner cortex in the left hemisphere (frontal, parietal, temporal, occipital) and 18 (out of 34) parcels of significantly thinner cortex (frontal, parietal, temporal, occipital) in the right hemisphere compared to monolinguals (Figure 5 and Supplemental Table 5).

### 3.5 Interaction of Language Background (Bilingual vs. Monolingual) and SES (Lower vs. Higher)

#### 3.5.1 Bilingual and Monolingual Groups with Lower and Higher SES demographic measures

As seen in Table 4, a one-way ANOVA of the demographic data showed that the four groups, Bilinguals of Lower SES, Monolinguals of Lower SES, Bilinguals of Higher SES and Monolinguals of Higher SES, did not differ in age, sex, and handedness. The ANOVA was significant for pubertal status for females, and t-tests revealed significant differences in the two Low-SES Groups (Bilinguals < Monolinguals), but not the two High-SES Groups. The ANOVA was also significant for nonverbal reasoning, but t-tests revealed no significant differences between the Bilingual and Monolingual Groups within the Low-SES or High-SES Groups. Note both variables were controlled for in the CT analysis. For 2×2 ANOVA results please see Supplemental Table 6.

**Table 4.** Demographics for the Lower-SES and the Higher-SES Bilingual Groups and Monolingual Groups.

|  | Lower-SES Groups |  | Higher-SES Groups |  | Statistics<br>One-way ANOVA |
| --- | --- | --- | --- | --- | --- |
|  | Bilingual | Monolingual | Bilingual | Monolingual |  |
| N | 322 | 318 | 223 | 247 | — |
| Age (years) | 9.83 (0.62) | 9.80 (0.60) | 9.80 (0.62) | 9.87 (0.61) | F(3)=0.69, n.s. |
| Pubertal Status - Female | 0.96 (1.23) | 1.28 (1.44) | 1.04 (1.28) | 0.84 (1.13) | F(3)=6.04, p < 0.05 |
| Pubertal Status - Male | 0.72 (0.89) | 0.82 (1.04) | 0.74 (0.88) | 0.76 (0.82) | F(3)=0.63, n.s. |
| Sex (M/F) | 168 / 154 | 159 / 159 | 117 / 106 | 137 / 110 | $\chi^2(3)=1.67$, n.s. |
| Handedness (R/Non-R) | 271 / 51 | 248 / 70 | 191 / 32 | 204 / 43 | $\chi^2(3)=6.57$, n.s. |
| SES: Household Income (USD) | 21,950 (12,500) | 19,850 (14,500) | 104,150 (49,000) | 111,750 (55,000) | F(3)=546.11, p < 0.05 |
| SES: Parental Education | 13.73 (3.52) | 13.73 (1.97) | 16.78 (2.62) | 16.56 (2.24) | F(3)=108.46, p < 0.05 |
| Nonverbal Reasoning (MR) | 8.98 (2.45) | 8.69 (2.57) | 10.26 (2.82) | 10.20 (2.59) | F(3)=26.88, p < 0.05 |

*Note:* Counts reported for group size, sex, and handedness. Averages and standard deviation reported for age, pubertal status, SES scores, nonverbal reasoning, vocabulary, language learning, inhibitory control, and visuospatial processing. Statistical tests: Student's t-test or chi-square, are reported in the right column. Pubertal status = Pubertal developmental scale; Sex: M=Male, F=Female; Handedness=Brief Edinburgh Handedness. SES Household Income=Parent Longitudinal Demographics Survey, in which household income bin was converted to income amount; SES Parental Education=Binned variable in which 13=High school graduate and 16=Associate degree: Occupational, Technical, or Vocational. Nonverbal; Reasoning=Matrix Reasoning (MR). Vocabulary=NIH Toolbox Picture Vocabulary Test (PVT) age-corrected standard score; Language Learning=Rey Auditory Verbal Learning Test (RAVLT) Learning score (Trial V-Trial I); Inhibitory Control=NIH Toolbox Flanker Task accuracy; Visuospatial Processing= Little Man Task percent correct. (n.s.=not significant, p ≥ 0.05).

#### 3.5.2 2×2 ANOVA for interaction of Language Background (Bilingual vs. Monolingual) and SES (Lower vs. Higher) on CT

Using vertex-wise analyses, we tested for an interaction of Language Background (Bilingual vs. Monolingual) x SES (Higher vs. Lower) on CT and found 11 clusters (Figure 6, Supplemental Table 6). Consistent with the prediction based on Brito and colleagues (Brito et al., 2018), three regions exhibited a pattern where bilinguals had greater CT than monolinguals in the Lower-SES Group, while there were no differences between bilinguals and monolinguals in the Higher-SES Group: the left inferior orbitofrontal gyrus, right fusiform gyrus, and right middle occipital gyrus. There were seven regions where the bilinguals again had greater CT than the monolinguals in the Lower-SES Group, but this time in the Higher-SES Group, bilinguals had less CT than monolinguals: right middle temporal gyrus, right precentral gyrus, right superior temporal gyrus, left inferior temporal gyrus, left fusiform gyrus, and left parahippocampal gyrus. Lastly, only one region, the left lingual gyrus, had a pattern where there were no differences between bilinguals and monolinguals in the Lower-SES Group, while in the Higher-SES Group bilinguals had less CT than monolinguals.

**Figure 6.**
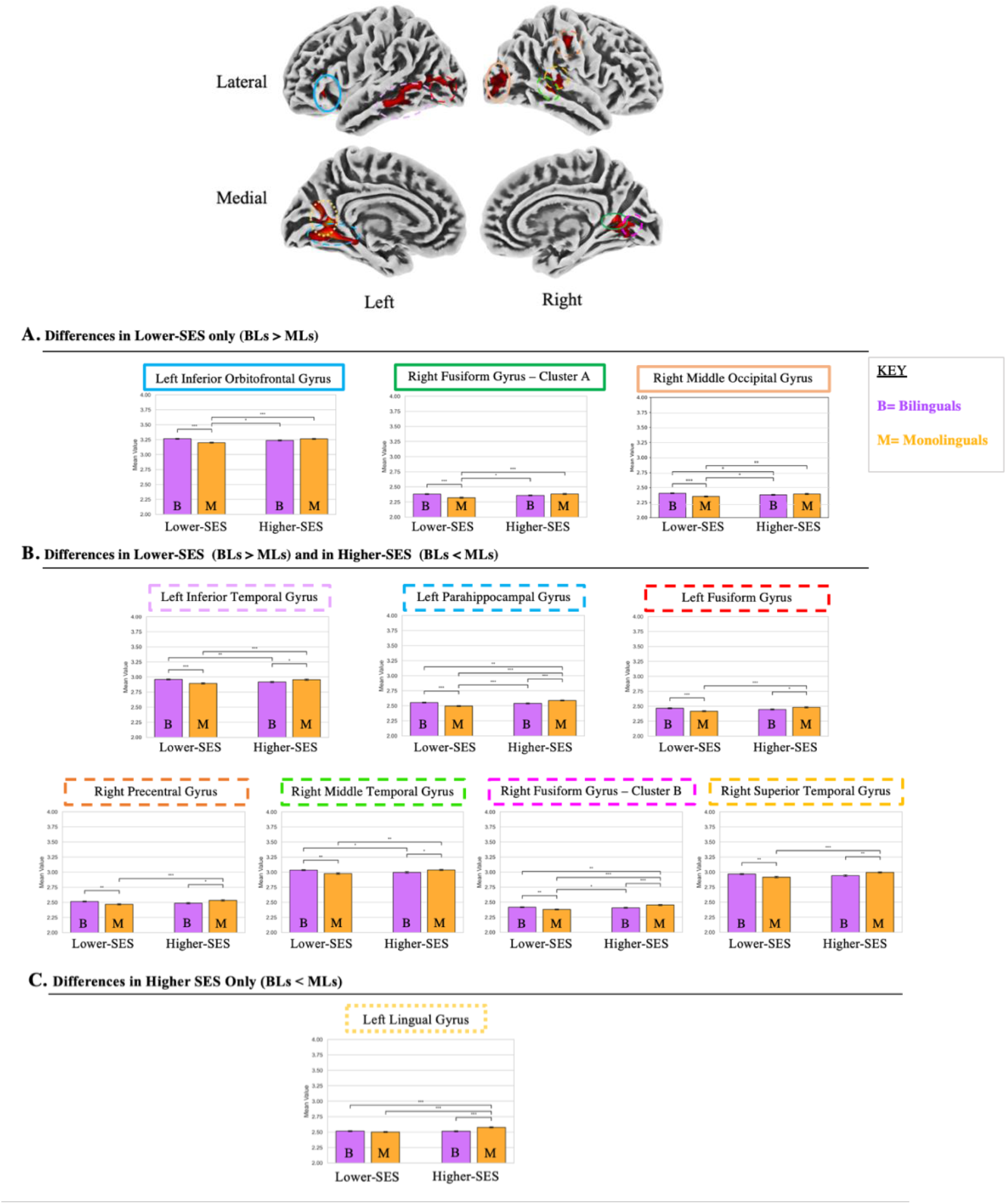
Language Background x SES Interaction on CT. *Note:* Bilinguals had more CT than monolinguals in the Lower-SES Group, while either (A) bilinguals and monolinguals in the Higher SES-Group did not differ in CT (solid circles/boxes), or (B) the bilinguals had less CT than monolinguals in the Higher-SES Group (dotted circles/boxes). (C) Bilinguals had less CT than monolinguals in the Higher-SES Group while there were no differences between bilinguals and monolinguals in the Lower-SES Group. 2 × 2 ANOVA analysis, with voxel-wise height threshold p<0.005, cluster-level extent threshold p<0.05, FDR corrected; p-values reported from post hoc student’s t-tests. Coordinates of peaks and subpeaks for each cluster are found in Supplementary Table 6.

## 4. Discussion

Few studies have investigated the effects of an early dual-language experience on cortical brain structure, and the results of these reports provide little convergence. In the present study, we compared a large group of early cultural Spanish-English bilinguals to English-speaking monolinguals. We conducted whole-brain, voxel/vertex-wise analyses of gray matter volume and cortical thickness, with CT as the main measure of interest and GMV for context with the predominant GMV literature. Between-group comparisons revealed that early bilinguals had relatively thicker cortex in regions within left inferior frontal, parahippocampal, occipital, but not parietal cortices, and within right superior temporal, occipital, but not frontal or parietal regions.

Relatively thinner cortex in early bilinguals was more extensive and involved regions in all lobes for both hemispheres. Differences identified between early bilinguals and monolinguals in prior studies have often been attributed to the bilinguals’ enhanced language and/or executive control skills involved in managing their two languages, but none tested whether the regions that distinguish early bilinguals from monolinguals are in fact related to performance on language or EC tasks in the same participants. Here, we conducted such correlations between CT and measures of language as well executive control skills but found no brain-behavioral relationships in those regions exhibiting between-group differences in CT to support the notion that either of these domains is tied to the adaptive structural changes borne out of the early dual-language experience, even when these skills differ between early bilinguals and monolinguals.

Given the frequent use of parcel-wise measures for CT data, we addressed the question of whether a parcel-wise approach leads to the same outcome for the comparison of early bilinguals and monolinguals by converting the vertex-wise data to average CT values in atlas-defined parcels. The results from a parcel-wise analyses were similar, albeit not identical, to those from the vertex-wise analyses when implemented with the Destrieux atlas, but less so when implemented with the Desikan-Killiany atlas, due to this atlas’ relatively fewer, larger parcels. Further, when conducting the same comparison of bilingual and monolingual groups on parcellated data provided by the ABCD Study Data Consortium, there again was divergence in the results from the Destrieux and Desikan-Killiany atlas, enough to give rise to alternate interpretations of the effect of an early dual-language experience on CT.

Lastly, given that neuroanatomical differences between bilinguals and monolinguals are affected by socioeconomic status (Brito and Noble et al., 2018), the current study controlled for household income and parental education by matching the early bilingual and monolingual groups on these measures and also entering them as a covariate of no interest in the analyses. However, we also directly tested for an interaction effect between language background (Bilingual vs. Monolingual) and SES (Higher vs Lower) on CT and found that amongst the Lower-SES Groups, there were numerous regions where bilinguals exhibited thicker cortex than monolinguals, while the Higher-SES Groups either did not differ, or differed in the opposite direction. These findings in CT are consistent with a pattern reported by Brito and Noble in 2018 for another neuroanatomical measure, underscoring that some brain structural differences between bilinguals and monolinguals are only identified if the groups are of lower SES. However, we also observed many regions that differed in the Bilingual and Monolingual Lower-SES Groups while the reverse effect manifested in the Bilingual and Monolingual Higher-SES Groups. These results reinforce the influential role of socioeconomic status across the spectrum on neuroanatomy.

### 4.1 Differences in GMV between early bilinguals and monolinguals

Most prior studies using a whole-brain, voxel-wise approach have reported GMD or GMV differences between early bilinguals and monolinguals, though the specific findings have varied. Mechelli identified more GMD in early bilinguals in the angular gyrus region of the left inferior parietal cortex, leading to interpretations that bilingualism modifies aspects of language. Given that overall vocabulary encompassing the two languages is larger in bilinguals than monolinguals (Allman, 2005; Bialystok, 2001; Pearson et al., 1993), it has been suggested that the acquisition and maintenance of a greater vocabulary might induce larger brain structures in left inferior parietal and inferior frontal cortices (Grogan et al, 2012; Green et al., 2007). However, Schug et al. (2022) found that early bilinguals had *less* GMV in the left supramarginal gyrus (Schug et al., 2022).

Specifically, Schug et al., (from here on considering the findings from children and adults combined) found more GMV in early bilinguals in left precentral gyrus and supplementary motor area, but the majority of findings of greater GMV in early bilinguals were in the right hemisphere, in pre- and postcentral gyrus extending into supramarginal gyrus, paracentral lobule extending to postcentral gyrus and inferior orbitofrontal gyrus (Schug et al., 2022). Notably, findings of less GMV were not only found in the left supramarginal gyrus, but also in left superior temporal gyrus extending to hippocampus, and numerous right hemisphere regions: Inferior frontal and orbital frontal, superior temporal extending into hippocampus, and occipital cortices. Less GMV was also identified in the left and right cerebellar hemispheres. The findings from Olulade et al., (2016) are consistent with these in terms of more GMV in right inferior frontal and inferior parietal cortices and in left and right precentral gyrus; and less GMV in left and right parahippocampal gyrus and cerebellum. Contrary to the focus on language in earlier work, both of these studies noted roles of right frontal and parietal cortices in executive control (Olulade et al., 2015; Schug et al., 2022), aligning more GMV in these regions with the theory that the bilingual’s constant need to monitor, select, and inhibit competing languages strengthens domain-general executive control systems (Green & Abutalebi, 2013). However, other studies found few (García-Pentón et al., 2019) or no group differences (Ressel et al., 2012) between early bilinguals and monolinguals.

This mix of findings was captured in a meta-analysis by (Danylkiv & Krafnick, 2020) which included all of the aforementioned GMD and GMV studies, except Schug et al., (2022) and included studies conducted in late bilinguals and reported no cortical differences between bilinguals and monolinguals. Inconsistent results may have been driven by varying methods and statistical thresholds, and small sample sizes. Mechelli et al., (2004) studied 25 early bilingual and 25 monolingual adults, Ressel et al., (2012) 22 early bilingual and 22 monolingual adults, Olulade et al., (2016) 15 early bilingual and 15 monolingual adults, García-Pentón et al., (2019) 14 early bilingual and 14 monolingual children, and Schug et al. (2022) 26 early bilingual and 42 monolingual adults, as well as 20 early bilingual and 34 monolingual children. The current study of 601 early bilingual and 608 monolingual children, and therefore has significantly more statistical power. In addition, the use of ABCD Study data afforded an opportunity to match for age, pubertal status, sex, handedness, SES, and nonverbal reasoning and to also control for these variables in the analyses.

In contrast to these prior studies, which reported several to no results, the current study revealed extensive differences in early bilinguals, relative to monolinguals, exhibiting more, as well as less GMV for all lobes in both hemispheres. Some of these results align with prior findings of more GMV in early bilinguals relative to monolinguals, with notable convergence in left supplementary motor area and right postcentral gyrus with Schug et al., (2022; adults and children combined), left middle occipital gyrus and right postcentral gyrus with Olulade et al., (2016), as well as right precuneus with García-Pentón et al., (2019). Also consistent with prior findings was less GMV in bilinguals compared to monolinguals in the left supramarginal gyrus, right superior temporal gyrus and occipital lobe with Schug et al., (2022), and left cuneus and right inferior temporal gyrus with Olulade et al., (2016), and bilateral cerebellum with Schug et al., (2022) and Olulade et al., (2016). In the context of the landmark study by Mechelli et al., (2004), we did not find more GMV in left inferior parietal cortex to implicate language, but early bilinguals did have more GMV in other language regions. Of note are left supplementary motor area/superior frontal gyrus, central to the production of spoken language (Vigneau et al., 2006), postcentral gyrus/superior parietal lobule, critical for written language (Menon & Desmond, 2001) and middle temporal gyrus, which provides lexical access to stored words (Sulpizio et al., 2020), together supporting language production and retrieval. At the same time, more GMV in right precuneus and less GMV in inferior parietal cortex could be indicative of adaptations involving executive control. Another notable result was more GMV in early bilinguals’ right occipital cortex together with precuneus, suggesting a possible role for visuospatial processing in dual-language users. In addition to these it is of interest that the current study and those by Schug et al., (2022) and Olulade et al., (2016), all identified less GMV in left and right cerebellum, tying the cerebellum into the potential mechanisms contemplated here.

Overall, findings from the current study show more extensive discrepancies in GMV than have been reported previously for whole-brain, voxel-wise studies in early bilinguals and provide evidence that an early dual-language experience substantially impacts GMV. This speaks to the importance of considering bilingualism as a factor in all GMV studies, given that early bilingualism presents with adaptations in GMV that are distributed across all cerebral and cerebellar lobes. Further, GMV differences in early bilinguals are expansive and do not point to one particular system that could account for the mechanism of experience-induced plasticity.

### 4.2. Differences in CT between early bilinguals and monolinguals

Of the two prior, whole-brain, vertex-wise studies of CT in early bilinguals, one revealed differences in two frontal regions (Klein et al., 2014) while the other had null results (Nguyen et al., 2023). These studies were conducted with 25 early bilingual and 22 monolingual adults, and 215 early bilingual and 145 monolingual children, respectively. The current study revealed much more widespread differences: Bilinguals exhibited relatively thicker cortex in regions of the left frontal, and the left and right temporal and occipital lobes.

Simultaneously, bilinguals showed extensive areas of relatively thinner cortex across all lobes and both hemispheres. Findings of thicker cortex in bilinguals in the left frontal lobe included a subpeak in the inferior orbitofrontal gyrus, which lies close to an inferior frontal gyrus region reported to have thicker cortex by Klein and colleagues (2014). However, our results did not reveal the same right inferior frontal regions with thinner cortex in the bilinguals relative to monolinguals as reported by Klein et al (2014). In addition, the current study revealed thicker cortex in left insula and parahippocampal gyrus, both involved in language, but also right middle temporal gyrus and left and right occipital cortex. These right hemisphere regions included early auditory and visual cortices and the large extent of thicker cortex in the left and right occipital lobes was unexpected, but consistent with the GMV results. On the other hand, regions of relatively thinner cortex in early bilinguals extended widely across the entire brain, encompassing those serving diverse functions. Regions involved in language, such as left inferior, middle and superior frontal gyri, supplementary motor area, supramarginal and precentral gyri were found to have thinner cortex in early bilinguals. At the same time, relatively thinner cortex in early bilinguals was also found in right superior frontal and precentral gyri, supplementary motor area and precuneus, which could be indicative of altered executive control systems in early bilinguals.

Taken together, the results for CT identified wide-spread differences in early bilinguals relative to monolinguals, with a predominance of thinner cortex. The results were similar to the GMV findings, with some variation in their location, as would be expected since they are not the same metric (GMV is a combination of CT and surface area). As noted above, the extent of findings makes it difficult to adjudicate between theories on the mechanisms by which the dual-language experience affects brain structure. To gain further insights, we next tested for correlations between CT and performance on language as well as EC measures in these regions of altered cortex.

### 4.3. CT Differences between bilinguals and monolinguals are not related to specific language or executive performance skills

In past work, neuroanatomical differences in early bilinguals have been tied to language and executive control by drawing on the literature, or by identifying correlations between brain structure and performance. For instance, in monolinguals, left and right inferior parietal GMD was found to correlate with vocabulary (Lee et al., 2007). In groups of mostly late bilinguals, left inferior parietal GMD was found to correlate with a combined language measure (Mechelli et al., 2004), and left inferior frontal GMD with a measure of lexical efficiency (Grogan et al., 2012). Here, we used measures of vocabulary and language learning and when comparing the groups on these measures, we found the early bilinguals to have a lower (English) vocabulary score than the monolinguals. This would be expected since bilinguals generally have a smaller vocabulary in each of their two languages, but an overall larger vocabulary when considering their combined vocabulary (Allman, 2005). On the other hand, the bilinguals scored higher on language learning than monolinguals did. Since these early cultural bilinguals learned their two languages as part of their family environment, it cannot be said that their stronger performance on this measure drove them to become bilinguals; instead, better memorization of word lists is likely the result of their bilingual experience. The test of inhibitory control was performed equally by the bilinguals and monolinguals, as expected given a prior behavioral study based on the ABCD Study participants (Dick et al., 2019), although counter to those that reported an advantage for inhibition in bilinguals (Bialystok et al., 2012; Bialystok & Werker, 2017). Lastly, the bilinguals performed relatively better on visuospatial processing, which we included post hoc given the relatively thicker cortex in left and right occipital lobes in bilinguals. These may result from the bilinguals’ reliance on visuospatial strategies to efficiently detect and respond to changes in their dual-language environment (Singh et al., 2024). Then we tested for correlations between CT (in all brain regions that differed in CT between the bilingual and monolingual groups) and performance on these measures but found no results.

Our study provides a much-needed investigation into brain-behavior associations that could explain what is driving the extensive neuroanatomical differences in early bilinguals. However, given the null findings, wide-spread alterations in CT likely reflect experience-dependent neuroplastic changes to support complex management of two languages that are not the result of any one specific enhanced skill. This null result fits with the position put forward by Blanco-Elorrieta and colleagues, who proposed that neuroanatomical differences observed in bilinguals may represent fine-tuned adaptations shaped by the varied and complex attentional, representational, and conversational demands of being bilingual, but not one specific skill (Blanco-Elorrieta & Caramazza, 2021; Blanco-Elorrieta, 2025). Also fitting with this theory is the fact that neuroanatomical differences did not coalesce around regions associated with one specific skill. It also explains the findings that bilinguals do not display heightened inhibitory control, yet still exhibit considerable divergence from monolinguals in their brain structure. Lastly, given the discrepancies in vocabulary, language learning, and visuospatial processing scores between the Bilingual and Monolingual Groups, we repeated the CT between-group comparison with these measures entered as a covariate of no interest to ensure they did not play a contributing role in the original results, but found the results to generally remain unaltered. Taken together, our findings do not support the notion that the adaptations in CT that follow an early dual-language experience are driven by a specific aspect of language or executive control skills.

### 4.4 Vertex-wise and parcel-wise analyses of same dataset does not produce the same results, especially when using a coarser atlas

Many reports on the ABCD Study data utilize the preprocessed parcellated averaged CT data provided by the ABCD Data Analysis, Informatics & Resource Center in tabulated format. The use of parcel values offers the convenience of time-efficient analyses and provides consistency among studies using the ABCD Study data. However, reducing vertex-wise information to average parcel values comes with a loss of information (Fürtjes et al., 2023) and studies reporting on between-group comparison using both vertex/voxel-wise and parcel-wise approaches simultaneously have documented this (Perlman et al., 2017; Wei et al., 2013, 2020). For example, Wei et al., (2013) found that Tai Chi practitioners exhibited thicker cortices in five clusters in left temporal, left occipital and right frontal lobes when taking a vertex-wise approach. However, the parcel-wise approach, whether using Destrieux or Desikan-Killiany atlases, detected no group differences. In the current study, we expanded the between-group comparisons of CT from our vertex-wise analysis to a parcel-wise analysis, to gain insights into the scope of such potential information loss. Further, we used both the Destrieux atlas (148 parcels) and the Desikan-Killiany atlas (68 parcels), given that atlas coarseness impacts the degree to which parcels maintain the information (Fürtjes et al., 2023). The first set of parcel-wise analyses were conducted by converting the vertex-wise data that were generated from the raw MRI data, and we then went on to conducted similar between-group analyses using the preprocessed ABCD Study data, providing two opportunities for side-by-side observations of CT differences in bilinguals with the Destrieux versus the Desikan-Killiany atlases.

We found that the conversion of vertex-wise data to parcel-wise data registered to the Destrieux atlas yielded results that largely resembled the vertex-wise findings, with all major regions identified in the vertex-wise analysis spatially aligned to corresponding parcels. The overall finding of a few clusters of thicker cortex in bilinguals together with numerous clusters of thinner cortex from the vertex-wise results was also observed in the Destrieux atlas parcellation results, with about 5% of parcels revealing relatively thicker, while 12% revealing relatively thinner cortex in bilinguals. However, despite general convergence, some regions identified in the vertex-wise analysis, e.g., thinner left supramarginal gyrus, were lost in the parcel-wise analysis using the Destrieux atlas, thus changing the interpretation for a region that is important to language.

While the Destrieux atlas generally captured many of the between-group differences identified in the vertex-wise analysis, the Desikan-Killiany atlas-based parcellation gave rise to a more spatially expanded version of these results due to the parcels being bigger and due to more parcels reporting significant results, especially for the Monolingual > Bilingual contrast. So, while 6% of parcels revealed thicker cortex in bilinguals than monolinguals under the Desikan-Killiany atlas (similar to the 5%, above), 24% of parcels indicated thinner cortex in bilinguals under the Desikan–Killiany atlas, double the 12% found under the Destrieux atlas. Parcels vary in size, so percentage values do not reflect the overall coverage, yet nevertheless provide some perspective. Specific examples of brain regions that would be interpreted differently under the Desikan–Killiany versus the Destrieux atlas results include the relatively thicker portions of the left occipital lobe and right primary auditory cortex within superior temporal lobe, which did not emerge from the Desikan–Killiany atlas. On the other hand, some regions appeared as significant in the Desikan-Killiany atlas, even though the Destrieux atlas (and the vertex-wise analysis) had not identified these, such as thinner cortex in early bilinguals in a large parcel representing the left inferior parietal cortex, and another representing the right superior temporal cortex. Notably the full length of the STG finding was likely driven by a difference in the temporal pole observed with the vertex-wise results (but absent in the Destrieux atlas results).

Turning to the ABCD-provided parcel-wise CT values analyzed for the comparison of the same Bilingual and Monolingual Groups, the results obtained via the two atlases demonstrated even more discrepancies. While the Destrieux atlas revealed 30% of parcels with relatively thicker and 49% of parcels with thinner cortex in bilinguals versus monolinguals, the Desikan-Killiany atlas yielded zero parcels of thicker and 52% of parcels with thinner cortex in bilinguals. Numerous regions identified as thicker in bilinguals under the Destrieux atlas were not detected under the Desikan-Killiany atlas, and others emerged with a flipped result, indicating thinner, rather than thicker CT in bilinguals. As such, use of the Desikan-Killiany atlas gave rise to alternate accounts of CT differences in early bilinguals.

These observations underscore the challenges associated with different analyses approaches, as articulated in a study of bilinguals which employed different neuroanatomical metrics and software packages (Claussenius-Kalman et al., 2020). Fürtjes et al., (2023), based on their quantitative comparison of several cortical atlases, recommended the use of vertex-wise analysis, or multiple fine-grained atlases for parcel-wise analyses. A prior study on CT in bilingual children from the ABCD Study was based on the Desikan-Killiany atlas (Vaughn et al., 2021). Despite some differences in participant selection and analysis methods, their results, using values provided by ABCD, are similar in spatial distribution to those reported in the current study for the same (Desikan-Killiany) atlas applied to the ABCD Study-provided data. They found one parcel (out of 68 or 1.5%) of thicker cortex (we found none) and 46 parcels (68%) with relatively thinner cortex in bilinguals (we found 35 or 52%). Yet the deterioration of fidelity of results when using the Desikan-Killiany over the Destrieux atlas observed in the current study suggests that the result by Vaughn et al., (2021) likely did not include all regions where early bilinguals differed from monolinguals, or included additional ones that are not true differences.

Taken together these additional analyses for between-group comparisons of CT reveal a relatively good, but not perfect, mapping from vertex-wise to parcel-wise results when utilizing the Destrieux atlas. As such, the benefits of time-efficient analyses and facilitation of cross-study comparisons have to be weighed against potential loss of some results. They also confirm low fidelity for results from the Desikan-Killiany atlas relative to the Destrieux atlas, as demonstrated for the same participants across two different analysis pipelines. It is noteworthy that CT differences in early bilinguals are especially extensive when considering other group comparisons in the broader literature. For example, between-group differences in CT for childhood conditions such as ADHD or developmental dyslexia (Schug & Eden, 2026) are more moderate, and these are at risk for being lost with the parcellated approach.

### 4.5 Socioeconomic Status Moderates Differences between Bilinguals and Monolingual in Cortical Thickness

Prior work by Brito and Noble reported that neuroanatomical differences in bilinguals were more prominent among adolescents from lower SES than from higher SES backgrounds. These findings were derived from a global average measure of cortical surface area (SA) (Brito & Noble, 2018). In the current study, our whole-brain vertex-wise approach resulted in multiple regions that exhibited a significant interaction between SES and Language Background on CT. For almost all regions that emerged from the interaction, early bilinguals had thicker cortex than monolinguals within the Lower-SES Groups but not the Higher-SES Groups, the latter mostly showing lower CT in bilinguals than monolinguals, or is some cases, no discrepancy. One possible interpretation is that dual-language exposure may serve as an enriching experience that amplifies experience-dependent neuroplasticity when overall environmental enrichment is lower. In higher-SES contexts, where cognitive and linguistic stimulation is more likely already abundant, the relative impact of bilingual experience on cortical structure may be attenuated or flipped. This position has been advanced by behavioral evidence suggesting that dual-language experience may moderate the effects of socioeconomic disadvantage on cognitive outcomes (Hartanto et al., 2019; Mezzacappa, 2004).

These two patterns can also be observed in some of the available performance measures. As can be seen from the 2×2 ANOVA (Supplementary Table 6), the dual-language experience provided a relative boost in visuospatial processing skills to the bilinguals in the Lower-SES Group, while there were no such differences between bilinguals and monolinguals in the Higher SES Groups. There was also a significant 2×2 interaction for inhibitory control, but no differences were determined post hoc between the Bilingual and Monolingual Groups at either SES level, indicating that this skill is not modulated by SES as might have been expected. For vocabulary, monolinguals in Higher-SES Groups had higher scores than the monolinguals, with no differences in the Lower-SES Groups. There was no interaction effect for language learning.

Taken together, the presence of this interaction effect on CT underscores the importance of considering SES in neuroanatomical studies of bilinguals. SES exerts widespread influences on cortical development and learning, and its interplay with a dual-language experience likely contributes to the heterogeneity of results across studies comparing bilinguals with monolinguals. Accounting for SES provides a more nuanced understanding of how social and linguistic environments jointly shape the developing brain.

## 5. Conclusion

Taken together, these findings from a large group of early cultural Spanish-English bilingual children matched to monolinguals, provide robust evidence that an early dual-language experience is associated with widespread differences in brain structure. However, the absence of brain-behavioral correlations suggest that such structural differences likely reflect broad experience-dependent plasticity rather than direct connections with specific language or executive control skills. Also, the use of both vertex-wise and parcel-wise approaches highlights the influence of methodological choice on the detection and interpretation of the neuroanatomical impact related to a dual-language experience. Lastly, there are multiple regions where SES interacts with language background to impact brain structure, emphasizing the need to consider the brain bases of bilingualism within broader sociocultural and environmental contexts.

## Data Availability Statement

All data files are available from the ABCD Study Data Repository from the NIMH Data Archive (https://nda.nih.gov/abcd).

## Acknowledgements

Data used in the preparation of this article were obtained from the Adolescent Brain Cognitive Development (ABCD) Study (https://abcdstudy.org), held in the NIMH Data Archive (NDA). This is a multisite, longitudinal study designed to recruit more than 10,000 children age 9–10 and follow them over 10 years into early adulthood. The ABCD Study® is supported by the National Institutes of Health and additional federal partners under award numbers U01DA041048, U01DA050989, U01DA051016, U01DA041022, U01DA0510 18, U01DA051037, U01DA050987, U01DA041174, U01DA041106, U01DA041117, U01DA041028, U01DA041134, U01DA050988, U01DA051039, U01DA041156, U01DA 041025, U01DA041120, U01DA051038, U01DA041148, U01DA041093, U01DA041089, U24DA041123, U24DA041147. A full list of supporters is available at https://abcdstudy.org/federal-partners.html.

The ABCD data repository grows and changes over time. The ABCD data used in this report came from the Fast Track data release. The raw data are available at https://nda.nih.gov/edit_collection.html?id=2573.

Instructions on how to create an NDA study are available at https://nda.nih.gov/nda/tutorials/creating_an_nda_study).

## Ethics Approval Statement

This study was performed in line with the principles of the Declaration of Helsinki. Parent consent and child assent was obtained by the ABCD Study (Garavan et al., 2018).

## Permission to reproduce material from other sources

N/A

