## Supplemental Tables for "Neuroanatomical differences in early bilingual and monolingual children"

**Supplemental Information**

**Supplemental Table 1. Locations of gray matter volume differences between the Bilingual and Monolingual Groups (voxel-wise analysis).**

| MNI Coordinates | | | | | | |
| --- | --- | --- | --- | --- | --- | --- |
| x | **y** | **z** | **Region** | **BA** | **Z** | **K_E_** |
| **Bilinguals > Monolinguals** | | | |  |  |  |
| **-6** | **15** | **66** | **Left Supplementary Motor Area** | **6** | **5.36** | **743** |
| -16 | -2 | 75 | Left Superior Frontal Gyrus | 6 | 4.75 |  |
| -9 | 2 | 72 | Left Supplementary Motor Area | 6 | 4.72 |  |
| **-57** | **-15** | **-16** | **Left Middle Temporal Gyrus** | **21** | **4.98** | **233** |
| -63 | -26 | -10 | Left Middle Temporal Gyrus | 21 | 4.6 |  |
| -70 | -30 | -9 | Left Middle Temporal Gyrus | 21 | 4.55 |  |
| **-16** | **-36** | **75** | **Left Postcentral Gyrus** | **1** | **4.85** | **314** |
| -16 | -45 | 70 | Left Precuneus | 7 | 4.61 |  |
| -24 | -44 | 74 | Left Superior Parietal Lobule | 7 | 4.23 |  |
| **-12** | **-102** | **-4** | **Left Calcarine Fissure** | **18** | **7.08** | **1931** |
| -18 | -102 | -10 | Left Inferior Occipital Gyrus | 18 | 6.74 |  |
| -28 | -98 | -4 | Left Middle Occipital Gyrus | 18 | 6.25 |  |
| **2** | **14** | **-14** | **Right Olfactory Gyrus** | **11** | **7.26** | **10578** |
| 30 | -15 | -21 | Right Hippocampus | 28 | 7.06 |  |
| 12 | -27 | -4 | Right Lingual Gyrus | 18 | 7.1 |  |
| **24** | **-39** | **74** | **Right Postcentral Gyrus** | **1** | **5.69** | **751** |
| 9 | -45 | 51 | Right Precuneus | 31 | 4.68 |  |
| 16 | -45 | 54 | Right Precuneus | 7 | 4.4 |  |
| **20** | **-100** | **-4** | **Right Calcarine Fissure** | **18** | **7.99** | **2008** |
| 30 | -96 | -2 | Right Inferior Occipital Gyrus | 18 | 7.3 |  |
| 38 | -93 | 2 | Right Middle Occipital Gyrus | 18 | 6.51 |  |
| **Bilinguals < Monolinguals** | | | |  |  |  |
| **-1** | **63** | **15** | **Left Medial Superior Frontal Gyrus** | **10** | **6.57** | **12062** |
| -2 | 60 | -24 | Left Gyrus Rectus | 11 | 6.63 |  |
| 12 | 62 | -26 | Right Superior Orbitofrontal Gyrus | 11 | 6.65 |  |
| **-22** | **20** | **-36** | **Left Superior Temporal Pole** | **38** | **6.17** | **688** |
| -18 | 6 | -34 | Left Superior Temporal Pole | 38 | 5.49 |  |
| -15 | 15 | -36 | Left Middle Temporal Pole | 38 | 4.51 |  |
| **-45** | **-34** | **30** | **Left Supramarginal Gyrus** | **40** | **4.87** | **425** |
| -40 | -38 | 24 | Left Superior Temporal Gyrus | 40 | 4.34 |  |
| -48 | -40 | 16 | Left Superior Temporal Gyrus | 22 | 4.01 |  |
| **-38** | **-69** | **9** | **Left Middle Occipital Gyrus** | **19** | **6.07** | **6765** |
| -6 | -90 | 38 | Left Cuneus | 19 | 5.8 |  |
| **-28** | **-74** | **-34** | **Left Cerebellum Crus I** | **-** | **7.86** | **16163** |
| 4 | -72 | -14 | Vermis XI | - | 6.23 |  |
| 22 | -76 | -32 | Right Cerebellum Crus I | - | 7.85 |  |
| **46** | **-39** | **21** | **Right Superior Temporal Gyrus** | **22** | **5.24** | **693** |
| 39 | -42 | 16 | Right Superior Temporal Gyrus | 22 | 4.67 |  |
| 42 | -44 | 28 | Right Angular Gyrus | 39 | 3.68 |  |
| **15** | **-48** | **-57** | **Right Cerebellum Lobule VIII** | **-** | **4.34** | **467** |
| 16 | -38 | -51 | Right Cerebellum Lobule IX | - | 4.04 |  |
| 20 | -46 | -44 | Right Cerebellum Lobule IX | - | 3.43 |  |
| **38** | **-51** | **-9** | **Right Fusiform Gyrus** | **37** | **5.1** | **389** |
| 44 | -39 | -12 | Right Inferior Temporal Gyrus | 37 | 4.26 |  |
| **39** | **-68** | **8** | **Right Middle Occipital Gyrus** | **19** | **4.71** | **367** |

*Note*: Report uses MNI Coordinates. Bold font indicates the peak coordinate of a cluster while italic font indicates the subpeaks within clusters.

**Supplemental Table 2**. **Locations of cortical thickness differences between the Bilingual and Monolingual Groups using the Destrieux atlas (values derived from vertex-wise results).**

| **Destrieux Parcel** | **F-stat** | **p-value** | **SS** | **Bilingual Mean** | **Monolingual Mean** |
| --- | --- | --- | --- | --- | --- |
| **Bilinguals > Monolinguals** |  |  |  |  |  |
| LEFT FRONTAL LOBE | | | | | |
| Anterior segment of the circular sulcus of the insula | 6.598 | p < 0.05 | 0.428 | 3.45 | 3.413 |
| LEFT PARIETAL LOBE |  |  |  |  |  |
| N/A |  |  |  |  |  |
| LEFT TEMPORAL LOBE |  |  |  |  |  |
| Parahippocampal gyrus (T5) | 6.511 | p < 0.05 | 0.206 | 2.925 | 2.894 |
| LEFT OCCIPITAL LOBE |  |  |  |  |  |
| Middle occipital sulcus and lunatus sulcus | 4.409 | p < 0.05 | 0.147 | 2.245 | 2.218 |
| Inferior occipital gyrus (O3) and sulcus | 7.079 | p < 0.05 | 0.223 | 2.382 | 2.322 |
| Occipital pole | 56.9 | p < 0.05 | 2.457 | 1.849 | 1.751 |
| RIGHT FRONTAL LOBE |  |  |  |  |  |
| N/A |  |  |  |  |  |
| RIGHT PARIETAL LOBE |  |  |  |  |  |
| N/A |  |  |  |  |  |
| RIGHT TEMPORAL |  |  |  |  |  |
| Anterior transverse temporal gyrus (Heschl) | 16.07 | p < 0.05 | 0.93 | 3.188 | 3.213 |
| RIGHT OCCIPITAL LOBE |  |  |  |  |  |
| Occipital pole | 43.286 | p < 0.05 | 1.262 | 2.94 | 2.964 |
| Inferior occipital gyrus (O3) and sulcus | 8.478 | p < 0.05 | 0.283 | 3.237 | 3.176 |
| **Bilinguals < Monolinguals** |  |  |  |  |  |
| LEFT FRONTAL LOBE |  |  |  |  |  |
| Transverse frontopolar gyri and sulci | 9.517 | p < 0.05 | 0.199 | 2.862 | 2.894 |
| Middle frontal gyrus (F2) | 17.38 | p < 0.05 | 0.3 | 2.943 | 2.977 |
| Superior frontal gyrus (F1) | 14.081 | p < 0.05 | 0.244 | 3.191 | 3.223 |
| Superior frontal sulcus | 11.447 | p < 0.05 | 0.134 | 2.879 | 2.904 |
| Subcentral gyrus (central operculum) and sulci | 8.577 | p < 0.05 | 0.218 | 2.872 | 2.9 |
| Central sulcus (Rolando’s fissure) | 10.993 | p < 0.05 | 0.194 | 2.174 | 2.201 |
| LEFT PARIETAL LOBE |  |  |  |  |  |
| Subparietal sulcus | 8.329 | p < 0.05 | 0.114 | 2.711 | 2.729 |
| Precuneus (medial part of P1) | 16.629 | p < 0.05 | 0.238 | 2.84 | 2.869 |
| Parieto-occipital sulcus | 8.112 | p < 0.05 | 0.145 | 2.549 | 2.57 |
| LEFT TEMPORAL LOBE |  |  |  |  |  |
| N/A |  |  |  |  |  |
| LEFT OCCIPITAL |  |  |  |  |  |
| Cuneus (O6) | 7.126 | p < 0.05 | 0.126 | 2.214 | 2.233 |
| Inferior occipital gyrus (O3) and sulcus | 5.599 | p < 0.05 | 0.213 | 2.553 | 2.522 |
| RIGHT FRONTAL LOBE |  |  |  |  |  |
| Transverse frontopolar gyri and sulci | 7.053 | p < 0.05 | 0.133 | 2.146 | 2.175 |
| Middle frontal gyrus (F2) | 7.199 | p < 0.05 | 0.136 | 2.628 | 2.587 |
| Superior frontal gyrus (F1) | 7.351 | p < 0.05 | 0.137 | 2.906 | 2.929 |
| Superior frontal sulcus | 8.666 | p < 0.05 | 0.12 | 2.254 | 2.28 |
| Central sulcus (Rolando’s fissure) | 10.248 | p < 0.05 | 0.186 | 1.934 | 1.863 |
| RIGHT PARIETAL LOBE |  |  |  |  |  |
| Precuneus (medial part of P1) | 7.065 | p < 0.05 | 0.096 | 2.861 | 2.892 |
| RIGHT TEMPORAL LOBE |  |  |  |  |  |
| N/A |  |  |  |  |  |
| RIGHT OCCIPITAL LOBE |  |  |  |  |  |
| Cuneus (O6) | 11.964 | p < 0.05 | 0.207 | 2.728 | 2.727 |

*Note:* F(1, 1,199). Significant findings equal FDR-corrected p-value < 0.05. SS= Sum of Squares.

**Supplemental Table 3**. **Locations of cortical thickness differences between the Bilingual and the Monolingual Groups using the Desikan-Killiany atlas (values derived from vertex-wise results).**

| **Desikan-Killiany Parcel** | **F-stat** | **p-value** | **SS** | **Bilingual Mean** | **Monolingual Mean** |
| --- | --- | --- | --- | --- | --- |
| **Bilinguals > Monolinguals** | |  |  |  |  |
| LEFT FRONTAL LOBE | |  |  |  |  |
| Lateral Orbitofrontal | 6.932 | p<0.05 | 0.086 | 3.032 | 3.024 |
| LEFT PARIETAL LOBE | |  |  |  |  |
| N/A |  |  |  |  |  |
| LEFT TEMPORAL LOBE | |  |  |  |  |
| Entorhinal | 4.699 | p<0.05 | 0.331 | 3.431 | 3.396 |
| LEFT OCCIPITAL LOBE | |  |  |  |  |
| N/A |  |  |  |  |  |
| RIGHT FRONTAL LOBE | |  |  |  |  |
| Lateral Orbitofrontal | 4.396 | p<0.05 | 0.051 | 2.975 | 2.973 |
| RIGHT PARIETAL LOBE | |  |  |  |  |
| N/A |  |  |  |  |  |
| RIGHT TEMPORAL LOBE | |  |  |  |  |
| N/A |  |  |  |  |  |
| RIGHT OCCIPITAL LOBE | |  |  |  |  |
| Lateral Occipital | 6.703 | p<0.05 | 0.099 | 2.25 | 2.203 |
| **Bilinguals < Monolinguals** | |  |  |  |  |
| LEFT FRONTAL LOBE | |  |  |  |  |
| Caudal Middle Frontal | 6.121 | p<0.05 | 0.09 | 2.921 | 2.947 |
| Superior Frontal | 5.503 | p<0.05 | 0.066 | 3.126 | 3.152 |
| Precentral | 5.943 | p<0.05 | 0.11 | 2.662 | 2.685 |
| LEFT PARIETAL LOBE | |  |  |  |  |
| Inferior Parietal | 3.971 | p<0.05 | 0.047 | 2.771 | 2.772 |
| Postcentral | 7.771 | p<0.05 | 0.15 | 2.397 | 2.42 |
| Paracentral | 4.437 | p<0.05 | 0.064 | 2.796 | 2.806 |
| Precuneus | 7.282 | p<0.05 | 0.075 | 2.733 | 2.75 |
| LEFT TEMPORAL LOBE | |  |  |  |  |
| Fusiform | 4.722 | p<0.05 | 0.05 | 2.818 | 2.836 |
| LEFT OCCIPITAL LOBE | |  |  |  |  |
| Cuneus | 10.189 | p<0.05 | 0.167 | 2.283 | 2.316 |
| RIGHT FRONTAL LOBE | |  |  |  |  |
| Caudal Middle Frontal | 6.55 | p<0.05 | 0.111 | 2.931 | 2.957 |
| Superior Frontal | 4.349 | p<0.05 | 0.054 | 3.131 | 3.15 |
| Precentral | 5.101 | p<0.05 | 0.094 | 2.638 | 2.654 |
| RIGHT PARIETAL LOBE | |  |  |  |  |
| Postcentral | 8.534 | p<0.05 | 0.167 | 2.396 | 2.417 |
| Paracentral | 8.35 | p<0.05 | 0.123 | 2.787 | 2.803 |
| RIGHT TEMPORAL LOBE | |  |  |  |  |
| Superior Temporal | 3.765 | p<0.05 | 0.181 | 3.235 | 3.257 |
| RIGHT OCCIPITAL LOBE | |  |  |  |  |
| Cuneus | 6.061 | p<0.05 | 0.101 | 2.281 | 2.31 |

*Note:* F(1, 1,199). Significant findings equal FDR-corrected p-value < 0.05. SS= Sum of Squares.

**Supplemental Table 4. Locations of cortical thickness differences between the Bilingual and Monolingual Groups using the Destrieux atlas (values derived from ABCD tabulated data).**

| Destrieux Parcel | F-stat | p-value | SS | Bilingual Mean | Monolingual Mean |
| --- | --- | --- | --- | --- | --- |
| **Bilinguals > Monolinguals** | | |  |  |  |
| LEFT FRONTAL LOBE | |  |  |  |  |
| Straight gyrus, Gyrus rectus | 8.84 | p<0.05 | 0.318 | 2.387 | 2.356 |
| Orbital sulci (H-shaped sulci) | 66.414 | p<0.05 | 2.286 | 2.2 | 2.1 |
| Subcallosal area, subcallosal gyrus | 94.5 | p<0.05 | 3.52 | 2.552 | 2.439 |
| Middle-anterior part of the cingulate gyrus and sulcus (aMCC) | 63.82 | p<0.05 | 1.294 | 2.518 | 2.45 |
| Middle frontal gyrus (F2) | 65.971 | p<0.05 | 1.416 | 2.92 | 2.848 |
| Middle frontal sulcus | 38.12 | p<0.05 | 1.614 | 2.904 | 2.817 |
| Subcentral gyrus (central operculum) and sulci | 37.421 | p<0.05 | 1.217 | 2.71 | 2.639 |
| Short insular gyri | 124.98 | p<0.05 | 15.586 | 2.287 | 2.033 |
| Anterior segment of the circular sulcus of the insula | 47.042 | p<0.05 | 4.092 | 2.429 | 2.296 |
| Inferior segment of the circular sulcus of the insula | 45.492 | p<0.05 | 2.932 | 2.945 | 2.844 |
| LEFT PARIETAL LOBE | |  |  |  |  |
| Superior parietal lobule (lateral part of P1) | 21.29 | p<0.05 | 1.529 | 3.359 | 3.275 |
| Intraparietal sulcus and transverse parietal sulci | 7.966 | p<0.05 | 0.386 | 2.517 | 2.479 |
| Angular gyrus | 24.017 | p<0.05 | 0.737 | 3.009 | 2.959 |
| Postcentral sulcus | 107.165 | p<0.05 | 4.805 | 2.688 | 2.552 |
| LEFT TEMPORAL LOBE | |  |  |  |  |
| Planum polare of the superior temporal gyrus | 149.661 | p<0.05 | 16.941 | 2.391 | 2.128 |
| Anterior transverse temporal gyrus (of Heschl) | 76.212 | p<0.05 | 5.863 | 2.5 | 2.352 |
| Transverse temporal sulcus | 16.715 | p<0.05 | 0.526 | 2.288 | 2.251 |
| Anterior transverse collateral sulcus | 71.13 | p<0.05 | 4.825 | 1.939 | 1.799 |
| Posterior transverse collateral sulcus | 51.147 | p<0.05 | 0.97 | 2.464 | 2.401 |
| Posterior ramus of the lateral sulcus (or fissure) | 60.879 | p<0.05 | 4.393 | 2.547 | 2.41 |
| LEFT OCCIPITAL LOBE | |  |  |  |  |
| Anterior occipital sulcus and preoccipital notch (temporo-occipital incisure) | 153.35 | p<0.05 | 33.807 | 2.32 | 1.962 |
| RIGHT FRONTAL LOBE | |  |  |  |  |
| Orbital sulci (H-shaped sulci) | 78.56 | p<0.05 | 3.093 | 2.175 | 2.06 |
| Suborbital sulcus (sulcus rostrales, supraorbital sulcus) | 7.676 | p<0.05 | 0.142 | 2.636 | 2.612 |
| Subcallosal area, subcallosal gyrus | 100.374 | p<0.05 | 4.159 | 2.524 | 2.395 |
| Middle-anterior part of the cingulate gyrus and sulcus (aMCC) | 86.655 | p<0.05 | 1.698 | 2.506 | 2.42 |
| Middle frontal gyrus (F2) | 74.25 | p<0.05 | 1.622 | 2.921 | 2.836 |
| Middle frontal sulcus | 33.655 | p<0.05 | 1.45 | 2.887 | 2.802 |
| Orbital part of the inferior frontal gyrus | 5.14 | p<0.05 | 0.071 | 2.903 | 2.887 |
| Subcentral gyrus (central operculum) and sulci | 27.267 | p<0.05 | 0.89 | 2.688 | 2.62 |
| Short insular gyri | 126.595 | p<0.05 | 14.592 | 2.296 | 2.056 |
| Anterior segment of the circular sulcus of the insula | 31.944 | p<0.05 | 2.83 | 2.444 | 2.337 |
| Inferior segment of the circular sulcus of the insula | 56.999 | p<0.05 | 3.376 | 2.963 | 2.837 |
| RIGHT PARIETAL LOBE | |  |  |  |  |
| Intraparietal sulcus and transverse parietal sulci | 4.999 | p<0.05 | 0.218 | 2.531 | 2.503 |
| Supramarginal gyrus | 19.95 | p<0.05 | 0.67 | 3.248 | 3.201 |
| Angular gyrus | 29.32 | p<0.05 | 0.914 | 3.016 | 2.957 |
| Postcentral sulcus | 118.653 | p<0.05 | 4.933 | 2.686 | 2.544 |
| RIGHT TEMPORAL LOBE | | |  |  |  |
| Planum polare of the superior temporal gyrus | 128.682 | p<0.05 | 14.259 | 2.451 | 2.209 |
| Anterior transverse temporal gyrus (of Heschl) | 66.693 | p<0.05 | 4.37 | 2.566 | 2.435 |
| Transverse temporal sulcus | 15.11 | p<0.05 | 0.439 | 2.357 | 2.32 |
| Anterior transverse collateral sulcus | 74.441 | p<0.05 | 4.876 | 1.917 | 1.774 |
| Posterior transverse collateral sulcus | 27.4 | p<0.05 | 0.595 | 2.427 | 2.371 |
| Posterior ramus of the lateral sulcus (or fissure) | 69.393 | p<0.05 | 5.371 | 2.52 | 2.374 |
| RIGHT OCCIPITAL LOBE | | |  |  |  |
| Lingual gyrus (O5) | 7.31 | p<0.05 | 0.182 | 2.508 | 2.488 |
| Superior occipital gyrus (O1) | 4.848 | p<0.05 | 0.081 | 2.362 | 2.348 |
| Anterior occipital sulcus and preoccipital notch (temporo-occipital incisure) | 147.213 | p<0.05 | 30.1 | 2.378 | 2.036 |
| **Bilinguals < Monolinguals** | | |  |  |  |
| LEFT FRONTAL LOBE | |  |  |  |  |
| Transverse frontopolar gyri and sulci | 24.033 | p<0.05 | 0.62 | 2.89 | 2.94 |
| Fronto-marginal gyrus (of Wernicke) and sulcus | 8.322 | p<0.05 | 0.21 | 2.57 | 2.6 |
| Medial orbital sulcus (olfactory sulcus) | 7.01 | p<0.05 | 0.18 | 2.6 | 2.62 |
| Anterior part of the cingulate gyrus and sulcus (ACC) | 7.085 | p<0.05 | 0.09 | 2.49 | 2.51 |
| Superior frontal gyrus (F1) | 56.18 | p<0.05 | 1.8 | 3.05 | 3.15 |
| Inferior frontal sulcus | 138.7 | p<0.05 | 11.74 | 3.27 | 3.49 |
| Triangular part of the inferior frontal gyrus | 5.8 | p<0.05 | 0.11 | 2.89 | 2.91 |
| Orbital part of the inferior frontal gyrus | 14.25 | p<0.05 | 0.21 | 2.92 | 2.96 |
| Horizontal ramus of the anterior segment of the lateral sulcus | 84.991 | p<0.05 | 3.24 | 2.88 | 2.99 |
| Vertical ramus of the anterior segment of the lateral sulcus | 56.043 | p<0.05 | 1.8 | 2.6 | 2.68 |
| Precentral gyrus | 109.239 | p<0.05 | 4.67 | 2.59 | 2.72 |
| Superior part of the precentral sulcus | 82.47 | p<0.05 | 2.2 | 2.92 | 3.02 |
| Central sulcus (Rolando’s fissure) | 83.28 | p<0.05 | 5.41 | 2.68 | 2.83 |
| Long insular gyrus and central sulcus of the insula | 119.56 | p<0.05 | 15.02 | 2.63 | 2.87 |
| Superior segment of the circular sulcus of the insula | 5.08 | p<0.05 | 0.28 | 3.32 | 3.36 |
| LEFT PARIETAL LOBE | |  |  |  |  |
| Sulcus intermedius primus (of Jensen) | 144.463 | p<0.05 | 15.71 | 3.05 | 3.29 |
| Postcentral gyrus | 161.3 | p<0.05 | 22.65 | 3.39 | 3.69 |
| Precuneus (medial part of P1) | 48.597 | p<0.05 | 1.17 | 2.51 | 2.57 |
| Subparietal sulcus | 62.37 | p<0.05 | 1.36 | 2.64 | 2.72 |
| Marginal branch of the cingulate sulcus | 156.8 | p<0.05 | 10.07 | 2.27 | 2.47 |
| Middle-posterior part of the cingulate gyrus and sulcus (pMCC) | 18.655 | p<0.05 | 0.44 | 2.65 | 2.68 |
| Posterior-dorsal part of the cingulate gyrus (dPCC) | 5.675 | p<0.05 | 0.5 | 2.54 | 2.58 |
| Posterior-ventral part of the cingulate gyrus (vPCC/isthmus) | 98.021 | p<0.05 | 1.64 | 2.33 | 2.41 |
| LEFT TEMPORAL LOBE | |  |  |  |  |
| Temporal pole | 20.099 | p<0.05 | 0.39 | 2.82 | 2.86 |
| Lateral aspect of the superior temporal gyrus | 159.96 | p<0.05 | 15.29 | 2.76 | 3 |
| Superior temporal sulcus (parallel sulcus) | 116.489 | p<0.05 | 7.67 | 2.64 | 2.81 |
| Middle temporal gyrus (T2) | 5.75 | p<0.05 | 0.2 | 2.94 | 2.97 |
| Inferior temporal gyrus (T3) | 15.543 | p<0.05 | 0.38 | 3.03 | 3.07 |
| Lateral occipito-temporal gyrus (fusiform gyrus, O4-T4) | 22.51 | p<0.05 | 0.95 | 3.11 | 3.17 |
| Lateral occipito-temporal sulcus | 177.777 | p<0.05 | 35.85 | 3 | 3.38 |
| Parahippocampal gyrus (T5) | 18 | p<0.05 | 0.39 | 2.6 | 2.62 |
| Medial occipito-temporal sulcus and lingual sulcus | 11.1 | p<0.05 | 0.19 | 2.65 | 2.67 |
| LEFT OCCIPITAL LOBE | |  |  |  |  |
| Calcarine sulcus | 94.777 | p<0.05 | 3.18 | 2.7 | 2.81 |
| Inferior occipital gyrus (O3) and sulcus | 7.185 | p<0.05 | 0.18 | 2.57 | 2.6 |
| Superior occipital sulcus and transverse occipital sulcus | 165.99 | p<0.05 | 11.01 | 2.45 | 2.66 |
| Occipital pole | 9.013 | p<0.05 | 0.39 | 2.95 | 3 |
| RIGHT FRONTAL LOBE | |  |  |  |  |
| Transverse frontopolar gyri and sulci | 31.65 | p<0.05 | 0.86 | 2.85 | 2.91 |
| Lateral orbital sulcus | 21.36 | p<0.05 | 0.38 | 2.12 | 2.15 |
| Anterior part of the cingulate gyrus and sulcus (ACC) | 4.512 | p<0.05 | 0.06 | 2.48 | 2.49 |
| Pericallosal sulcus (S of corpus callosum) | 10.322 | p<0.05 | 0.16 | 2.58 | 2.6 |
| Superior frontal gyrus (F1) | 27.558 | p<0.05 | 0.82 | 3.09 | 3.15 |
| Inferior frontal sulcus | 128.859 | p<0.05 | 9.3 | 3.18 | 3.38 |
| Triangular part of the inferior frontal gyrus | 7.926 | p<0.05 | 0.13 | 2.91 | 2.93 |
| Horizontal ramus of the anterior segment of the lateral sulcus | 64.65 | p<0.05 | 2.66 | 2.9 | 3.01 |
| Vertical ramus of the anterior segment of the lateral sulcus | 45.82 | p<0.05 | 1.75 | 2.61 | 2.7 |
| Precentral gyrus | 129.532 | p<0.05 | 5.19 | 2.67 | 2.81 |
| Inferior part of the precentral sulcus | 13 | p<0.05 | 0.47 | 3.02 | 3.07 |
| Superior part of the precentral sulcus | 32.88 | p<0.05 | 0.89 | 2.88 | 2.94 |
| Central sulcus (Rolando’s fissure) | 58.172 | p<0.05 | 3.08 | 2.69 | 2.81 |
| Long insular gyrus and central sulcus of the insula | 136.893 | p<0.05 | 17.93 | 2.69 | 2.95 |
| Superior segment of the circular sulcus of the insula | 11.496 | p<0.05 | 0.52 | 3.31 | 3.35 |
| RIGHT PARIETAL LOBE | |  |  |  |  |
| Paracentral lobule and sulcus | 17.275 | p<0.05 | 0.64 | 2.67 | 2.72 |
| Sulcus intermedius primus (of Jensen) | 154.854 | p<0.05 | 14.98 | 3.04 | 3.28 |
| Postcentral gyrus | 125.155 | p<0.05 | 13.6 | 3.28 | 3.52 |
| Precuneus (medial part of P1) | 54.498 | p<0.05 | 1.3 | 2.52 | 2.59 |
| Subparietal sulcus | 25.31 | p<0.05 | 0.51 | 2.72 | 2.77 |
| Marginal branch of the cingulate sulcus | 144.687 | p<0.05 | 10.16 | 2.28 | 2.48 |
| Middle-posterior part of the cingulate gyrus and sulcus (pMCC) | 14.593 | p<0.05 | 0.3 | 2.62 | 2.65 |
| Posterior-dorsal part of the cingulate gyrus (dPCC) | 25.162 | p<0.05 | 0.82 | 2.47 | 2.53 |
| Posterior-ventral part of the cingulate gyrus (vPCC/isthmus) | 100.74 | p<0.05 | 1.55 | 2.33 | 2.41 |
| RIGHT TEMPORAL LOBE | | |  |  |  |
| Temporal pole | 12.25 | p<0.05 | 0.23 | 2.82 | 2.85 |
| Lateral aspect of the superior temporal gyrus | 162.81 | p<0.05 | 14.54 | 2.8 | 3.04 |
| Superior temporal sulcus (parallel sulcus) | 85.145 | p<0.05 | 4.87 | 2.67 | 2.81 |
| Planum temporale | 6.746 | p<0.05 | 0.27 | 3.13 | 3.17 |
| Middle temporal gyrus (T2) | 8.916 | p<0.05 | 0.27 | 2.96 | 2.99 |
| Inferior temporal gyrus (T3) | 29.958 | p<0.05 | 0.66 | 3.05 | 3.1 |
| Lateral occipito-temporal gyrus (fusiform gyrus, O4-T4) | 11.099 | p<0.05 | 0.53 | 3.07 | 3.11 |
| Lateral occipito-temporal sulcus | 180.3 | p<0.05 | 34.37 | 3.05 | 3.42 |
| Parahippocampal gyrus (T5) | 9.625 | p<0.05 | 0.23 | 2.61 | 2.64 |
| Medial occipito-temporal sulcus and lingual sulcus | 16.554 | p<0.05 | 0.34 | 2.62 | 2.65 |
| RIGHT OCCIPITAL LOBE | | |  |  |  |
| Calcarine sulcus | 62.02 | p<0.05 | 2.03 | 2.7 | 2.79 |
| Superior occipital sulcus and transverse occipital sulcus | 182.766 | p<0.05 | 11.19 | 2.51 | 2.71 |
| Occipital pole | 6.361 | p<0.05 | 0.32 | 2.92 | 2.96 |

*Note:* F(1, 1,199). Significant findings equal FDR-corrected p-value < 0.05. SS= Sum of Squares.

**Supplemental Table 5.** **Locations of cortical thickness differences between the Bilingual and Monolingual Groups using the Desikan-Killiany atlas (values derived from ABCD tabulated data)**

| Desikan-Killiany Parcel | F-stat | p-value | SS | Bilingual Mean | Monolingual Mean |
| --- | --- | --- | --- | --- | --- |
| **Bilinguals > Monolinguals** | | |  |  |  |
| LEFT FRONTAL LOBE | |  |  |  |  |
| N/A |  |  |  |  |  |
| LEFT PARIETAL LOBE | |  |  |  |  |
| N/A |  |  |  |  |  |
| LEFT TEMPORAL LOBE | |  |  |  |  |
| N/A |  |  |  |  |  |
| LEFT OCCIPITAL LOBE | |  |  |  |  |
| N/A |  |  |  |  |  |
| **Bilinguals < Monolinguals** | | |  |  |  |
| LEFT FRONTAL LOBE | |  |  |  |  |
| Rostral Anterior Cingulate | 22.626 | p < 0.05 | 0.689 | 2.939 | 2.874 |
| Pars Opercularis | 8.94 | p < 0.05 | 0.127 | 2.871 | 2.848 |
| Frontal pole | 5.494 | p < 0.05 | 0.338 | 3.101 | 3.052 |
| Caudal Middle Frontal | 4.328 | p < 0.05 | 0.074 | 2.848 | 2.833 |
| Isthmus Cingulate | 8.7 | p < 0.05 | 0.163 | 2.465 | 2.44 |
| LEFT PARIETAL LOBE | |  |  |  |  |
| Superior Parietal | 7.145 | p < 0.05 | 0.082 | 2.501 | 2.481 |
| Inferior Parietal | 21.018 | p < 0.05 | 0.285 | 2.729 | 2.695 |
| Supramarginal | 7.301 | p < 0.05 | 0.126 | 2.816 | 2.8 |
| Postcentral | 18.058 | p < 0.05 | 0.292 | 2.33 | 2.301 |
| Precuneus | 14.501 | p < 0.05 | 0.148 | 2.672 | 2.649 |
| LEFT TEMPORAL LOBE | |  |  |  |  |
| Superior Temporal | 23.671 | p < 0.05 | 0.459 | 3.052 | 3.006 |
| Transverse Temporal | 9.255 | p < 0.05 | 0.28 | 2.757 | 2.725 |
| Middle Temporal | 9.283 | p < 0.05 | 0.228 | 3.071 | 3.048 |
| Inferior Temporal | 17.325 | p < 0.05 | 0.299 | 2.969 | 2.946 |
| LEFT OCCIPITAL LOBE | |  |  |  |  |
| Cuneus | 22.321 | p < 0.05 | 0.354 | 2.109 | 2.075 |
| Lateral Occipital | 18.56 | p < 0.05 | 0.262 | 2.308 | 2.275 |
| Pericalcarine | 17.72 | p < 0.05 | 0.337 | 1.841 | 1.806 |
| RIGHT FRONTAL LOBE | |  |  |  |  |
| Rostral Anterior Cingulate | 10.128 | p < 0.05 | 0.347 | 2.931 | 2.893 |
| Pars Opercularis | 7.838 | p < 0.05 | 0.11 | 2.873 | 2.846 |
| Insula | 10.526 | p < 0.05 | 0.206 | 3.197 | 3.162 |
| Isthmus Cingulate | 11.636 | p < 0.05 | 0.221 | 2.464 | 2.44 |
| RIGHT PARIETAL LOBE | |  |  |  |  |
| Superior Parietal | 5.035 | p < 0.05 | 0.058 | 2.504 | 2.489 |
| Inferior Parietal | 24.572 | p < 0.05 | 0.324 | 2.76 | 2.722 |
| Supramarginal | 9.318 | p < 0.05 | 0.154 | 2.813 | 2.789 |
| Postcentral | 7.475 | p < 0.05 | 0.151 | 2.309 | 2.289 |
| Precuneus | 5.555 | p < 0.05 | 0.053 | 2.678 | 2.664 |
| RIGHT TEMPORAL | |  |  |  |  |
| Superior Temporal | 20.106 | p < 0.05 | 0.376 | 3.075 | 3.035 |
| Transverse Temporal | 5.692 | p < 0.05 | 0.174 | 2.768 | 2.753 |
| Middle Temporal | 9.055 | p < 0.05 | 0.2 | 3.084 | 3.063 |
| Inferior Temporal | 14.634 | p < 0.05 | 0.246 | 2.991 | 2.962 |
| Temporal Pole | 4.383 | p < 0.05 | 0.3 | 3.77 | 3.729 |
| RIGHT OCCIPITAL LOBE | | |  |  |  |
| Lateral Occipital | 13.479 | p < 0.05 | 0.199 | 2.375 | 2.347 |
| Pericalcarine | 15.501 | p < 0.05 | 0.317 | 1.836 | 1.807 |
| Cuneus | 14.367 | p < 0.05 | 0.229 | 2.14 | 2.109 |
| Lingual | 3.018 | p < 0.05 | 0.041 | 2.227 | 2.218 |

*Note:* F(1, 1,199). Significant findings equal FDR-corrected p-value < 0.05. SS= Sum of Squares.

**Table 6. 2x2 ANOVA of demographic and performance measures for the Lower-SES and Higher-SES Bilingual and Monolingual Groups.**

|  | **Lower-SES** | | **Higher-SES** | | **ANOVA**  **inter-action** | **Post hoc test**  **(within SES level)** | |
| --- | --- | --- | --- | --- | --- | --- | --- |
|  | **Bilinguals** | **Monolinguals** | **Bilinguals** | **Monolinguals** |  | **Lower-SES** | **Higher-SES** |
| **N** | 322 | 318 | 223 | 247 | n.s. | — | — |
| **Age (years)** | 9.83 ± 0.62 | 9.80 ± 0.60 | 9.80 ± 0.62 | 9.87 ± 0.61 | n.s. | — | — |
| **Pubertal Status – Female** | 0.96 ± 1.23 | 1.28 ± 1.44 | 1.04 ± 1.28 | 0.84 ± 1.13 | F= 11.27 | BL<ML | BL=ML |
| **Pubertal Status – Male** | 0.72 ± 0.89 | 0.82 ± 1.04 | 0.74 ± 0.88 | 0.76 ± 0.82 | n.s. | — | — |
| **Sex (M / F)** | 168 / 154 | 159 / 159 | 117 / 106 | 137 / 110 | n.s. | — | — |
| **Handedness (R/Non-R)** | 271 / 51 | 248 / 70 | 191 / 32 | 204 / 43 | n.s. | — | — |
| **SES: Household Income (USD)** | 21,950 ± 12,500 | 19,850 ± 14,500 | 104,150 ± 49,000 | 111,750 ± 55,000 | F=5.03 | BL=ML | BL=ML |
| **SES: Parental Education** | 13.73 ± 3.52 | 13.73 ± 1.97 | 16.78 ± 2.62 | 16.56 ± 2.24 | n.s. |  |  |
| **Nonverbal Reasoning** | 8.98 ± 2.45 | 8.69 ± 2.57 | 10.26 ± 2.82 | 10.20 ± 2.59 | n.s. | — | — |
| **Vocabulary** | 98.15 ± 12.80 | 99.31 ± 13.39 | 105.49 ± 14.48 | 110.89 ± 15.72 | F= 6.21 | BL=ML | BL<ML |
| **Language Learning** | 6.27 ± 2.77 | 5.81 ± 2.69 | 6.35 ± 2.56 | 6.27 ± 2.77 | n.s. | — | — |
| **Inhibitory Control** | 95.02 ± 13.03 | 92.51 ± 12.48 | 95.40 ± 13.66 | 96.60 ± 12.83 | F= 5.54 | BL=ML | BL=ML |
| **Visuospatial Processing** | 0.57± 0.17 | 0.53 ± 0.15 | 0.61 ± 0.15 | 0.62 ± 0.18 | F=8.62 | BL>ML | BL=ML |

*Note:* Counts reported for group size, sex, and handedness. Averages and standard deviation (M±SD) reported for age, pubertal status, SES scores, nonverbal reasoning, vocabulary, language learning, inhibitory control, and visuospatial processing. BL=Bilingual, ML=Monolingual. Statistical tests: 2x2 ANOVAs for Bilingualism × SES interaction F(1, 1,106). Post hoc Tukey Test to identify significant between-group difference within the Lower-SES and the Higher SES Groups. Significant findings equal p-value < 0.05.

**Supplemental Table 7. Locations of interaction effects of Language Background (Bilingual vs. Monolingual) and SES (Lower vs. Higher) on cortical thickness (voxel-wise analysis).**

| **MNI Coordinates** | | | | | | |
| --- | --- | --- | --- | --- | --- | --- |
| **x** | **y** | **z** | **Region** | **BA** | **Z** | **KE** |
| Differences in Lower SES only (BLs > MLs) | | | | | | |
| **-34** | **44** | **-17** | **Left Inferior Orbitofrontal Gyrus** | **47** | **4.46** | **115** |
| **20** | **-32** | **-16** | **Right Fusiform Gyrus - Cluster A** | **36** | **4.51** | **416** |
| 10 | -43 | -7 | Right Parahippocampal Gyrus | 36 | 2.78 |  |
| 37 | -84 | **1** | **Right Middle Occipital Gyrus** | **18** | **4.48** | **565** |
| 33 | -81 | 8 | Right Middle Occipital Gyrus | 19 | 3.9 |  |
| Differences in Lower SES (BLs > MLs) and in Higher SES (BLs < MLs) | | | | | | |
| **-55** | **-21** | **-27** | **Left Inferior Temporal Gyrus** | 20 | 3.93 | 291 |
| -61 | -17 | -38 | Left Inferior Temporal Gyrus | 20 | 3.44 |  |
| -64 | -2 | -36 | Left Inferior Temporal Gyrus | 38 | 3.44 |  |
| **-14** | **-29** | **-16** | **Left Parahippocampal Gyrus** | 36 | 4.87 | 533 |
| -14 | -36 | -26 | Left Parahippocampal Gyrus | 37 | 4.34 |  |
| -16 | -22 | -30 | Left Parahippocampal Gyrus | 36 | 4.09 |  |
| Differences in Higher SES only (BLs < MLs) | | | | | | |
| **-43** | **-54** | **-23** | **Left Fusiform Gyrus** | **37** | **3.48** | **168** |
| **42** | **-5** | **28** | **Right Precentral Gyrus** | **6** | **3.99** | **177** |
| **57** | **-2** | **-16** | **Right Middle Temporal Gyrus** | **22** | **4.1** | **106** |
| **44** | **-50** | **-19** | **Right Fusiform Gyrus -Cluster B** | **37** | **4.09** | **330** |
| 39 | -54 | 6 | Right Middle Temporal Gyrus | 37 | 3.19 |  |
| 38 | -48 | 0 | Right Fusiform Gyrus | 21 | 3.14 |  |
| **56** | **2** | **-7** | **Right Superior Temporal Gyrus** | **22** | **4** | **210** |
| 56 | -10 | 5 | Right Superior Temporal Gyrus | 41 | 3.35 |  |
| 67 | -2 | 3 | Right Superior Temporal Gyrus | 22 | 3.03 |  |

*Note:* Report uses MNI Coordinates. Bold font indicates the peak coordinate of a cluster while italic font indicates the subpeaks within clusters.
